# Mycobacteriophage TM4 requires XylR for successful infection in *Mycobacterium smegmatis* mc^2^155

**DOI:** 10.64898/2026.05.17.725743

**Authors:** Annesha Adhikari, Tejan Lodhiya, Subhashree Nayak, Rohit Satardekar, Raju Mukherjee

## Abstract

Mycobacteriophages are viruses that infect mycobacteria, including *Mycobacterium tuberculosis*, and have emerged as promising alternatives to antibiotics in the face of increasing antimicrobial resistance. However, evolution of phage-resistance remains a major challenge to the clinical implementation of phage therapy. We describe a TM4 phage resistant mutant in *M. smegmatis* harboring an in-frame deletion in the *xylR*, transcriptional regulator implicated in lipid metabolism, and cell envelope homeostasis. Using spontaneous mutagenesis, transcriptomics and biochemical approaches, we identify a previously uncharacterized resistance mechanism mediated by cell-envelope remodeling that impedes productive phage infection. The *xylR* mutation disrupted phage DNA injection through enhanced recruitment of lipooligosaccharides to the cell surface, without inhibiting phage adsorption, genome replication, and virion assembly. Remodeling of the cell envelope was further enhanced by the induction of lipooligosaccharide biosynthesis upon TM4 infection, however, the phenotype can be reverted through chemical treatment, restoring phage sensitivity. Our study expands the paradigm of innate mechanisms underlying broad-spectrum phage resistance in mycobacteria.

## INTRODUCTION

Viruses infecting bacteria, known as bacteriophages, are the most abundant biological entities on Earth with remarkable genetic diversity and potential for use as alternate to antibiotics^1–3^. Phage infection begins with recognition of specific receptors on the bacterial cell surface, followed by injection of the viral genome inside the bacteria, replication within the host cell, and, ultimately, lysis of the bacterium to release numerous progeny phages. As bacteria are constantly exposed to phage predation in nature, they have evolved multiple defense strategies^2–4^. These include adaptive immune responses such as CRISPR–Cas, as well as innate immune systems like restriction–modification, toxin–antitoxin systems, BREX, and abortive infections. In turn, phages also evolve to counter the coevolutionary arms race that shapes viral host range and improves its therapeutic potential^5–9^. Prevention of phage adsorption through modification or loss of bacterial surface receptors constitutes a primary level of bacterial defense^10,11^.

In *Mycobacterium smegmatis*, resistance can arise not only from mutations affecting surface glycoconjugates such as glycopeptidolipids and trehalose polyphleates^12,13^, but also from intracellular mechanisms. These include overexpression of exonucleases such as multicopy phage resistance (Mpr)^14,15^, as well as mutations in the regulatory gene *lsr2*^16^. Mutations affecting receptor biosynthesis, extracellular matrix production, or essential intracellular host factors required for phage replication can all confer resistance^9^. Mycobacterial species, which includes the pathogens *M. tuberculosis* and *M. leprae*, encode both adaptive and innate anti-phage defense systems^17^. However, despite these recent findings, our understanding of molecular mechanisms of anti-phage resistance employed by mycobacteria remains poorly understood. A complete characterization of these mechanisms is a prerequisite, should mycobacteriophages be offered as adjunctive therapy for mycobacterial infection.

D-xylose is the most abundant fermentable pentose and acts as an important carbon and energy source for many bacteria^18–22^. Its metabolism is primarily regulated by *xylR*, a global transcription regulator that controls the expression of xylose uptake and catabolism^22–24^. In most bacterial species, *xylR* functions as a negative regulator in the absence of D-xylose, repressing the target genes expression^22^. Binding of D-xylose induces conformational changes in XylR that modulate its regulatory activity. Notably, the regulatory role of XylR can differ among species. In *Caulobacter crescentus*, D-xylose relieves XylR-mediated repression^22^, whereas in *E. coli*, XylR acts as a positive regulator^23,24^. In *M. smegmatis*, xylose has been detected in surface-exposed materials of the cell envelope^22–24^, suggesting a potential structural and regulatory significance. Studies reported that XylR acts as a negative regulator by repressing 166 involved in lipid synthesis and metabolism. Activation of XylR leads to alterations in lipid metabolism and multiple phenotypic alterations, such as cell size, colony morphology, biofilm formation, aggregation, and drug resistance^25^. Among the surface exposed glycoconjugates, lipooligosaccharides (LOS) contribute to sliding motility, biofilm formation, and macrophage infection^26–29^. First identified in *M. kansasii*^30^ and *M. smegmatis*^31^, LOS have since been reported in multiple other species, including “*M. canettii*” and related members of the *M. tuberculosis* complex^26,32–34^. Importantly, upregulation of the 11 gene cluster encompassing MSMEG_4727 has recently been associated with enhanced phage resistance in *M. smegmatis*^35^. In *M. smegmatis*, a 15-gene cluster was attributed for biosynthesis of the LOS glycoconjugate which comprise of a poly-*O*-acylated trehalose core decorated with mono- or oligosaccharide units^33^. An important characteristic of these molecules is presence of polymethyl-branched fatty acids, synthesized by a Mas-type polyketide synthase that utilizes methylmalonyl-CoA to generate branched aliphatic chains^36–38^. Despite their distinctive structural features, LOS biosynthesis remains incompletely understood, with only a limited number of genes experimentally validated in the pathway^26,39^.

In this study, we demonstrate that infection by Mycobacteriophage TM4 selects for mutations in the *xylR* regulator, leading to the de-repression of the LOS synthesis gene island (MSMEG_4727–4737). This activation confers cross resistance to mycobacterial phages due to the production of additional LOS. Our findings show a previously uncharacterized XylR-mediated signaling pathway that links metabolic regulation to an induced innate immunity against phages and provide new insights into the co-evolutionary dynamics between mycobacterial hosts and their phages.

## RESULTS

### Isolation of a spontaneous mutant of *M. smegmatis* mc^2^155 resistant to TM4 Mycobacteriophage

To decipher new mechanisms of phage resistance in *Mycobacterium smegmatis*, we employed a spontaneous mutagenesis approach using the mycobacteriophage MycoMarT7 (TM4). MycoMarT7 is an engineered derivative of the cluster K mycobacteriophage TM4 and was chosen for its ability to efficiently lyse *M. smegmatis* at 30 °C. Cells were challenged with MycoMarT7 at a multiplicity of infection (MOI) of 1:5 and after various time intervals, aliquots of cells were plated on 7H10 plates, and incubated at 30 °C, that is the permissive for MycoMar T7 phage replication and lysis. Following infection, few colonies exhibiting resistance to MycoMarT7 were isolated. For ensuring the stability of the resistant phenotype, the mutants were subjected to five sequential rounds of high-MOI infection with MycoMarT7. After each round, cultures were plated and examined for plaque formation. The colony that maintained total resistance throughout all passages, exhibiting no lytic zones at the highest titre, was subsequently designated as the *phage-resistant mutant* (PRM) (Figure 1a).

**Figure 1:**
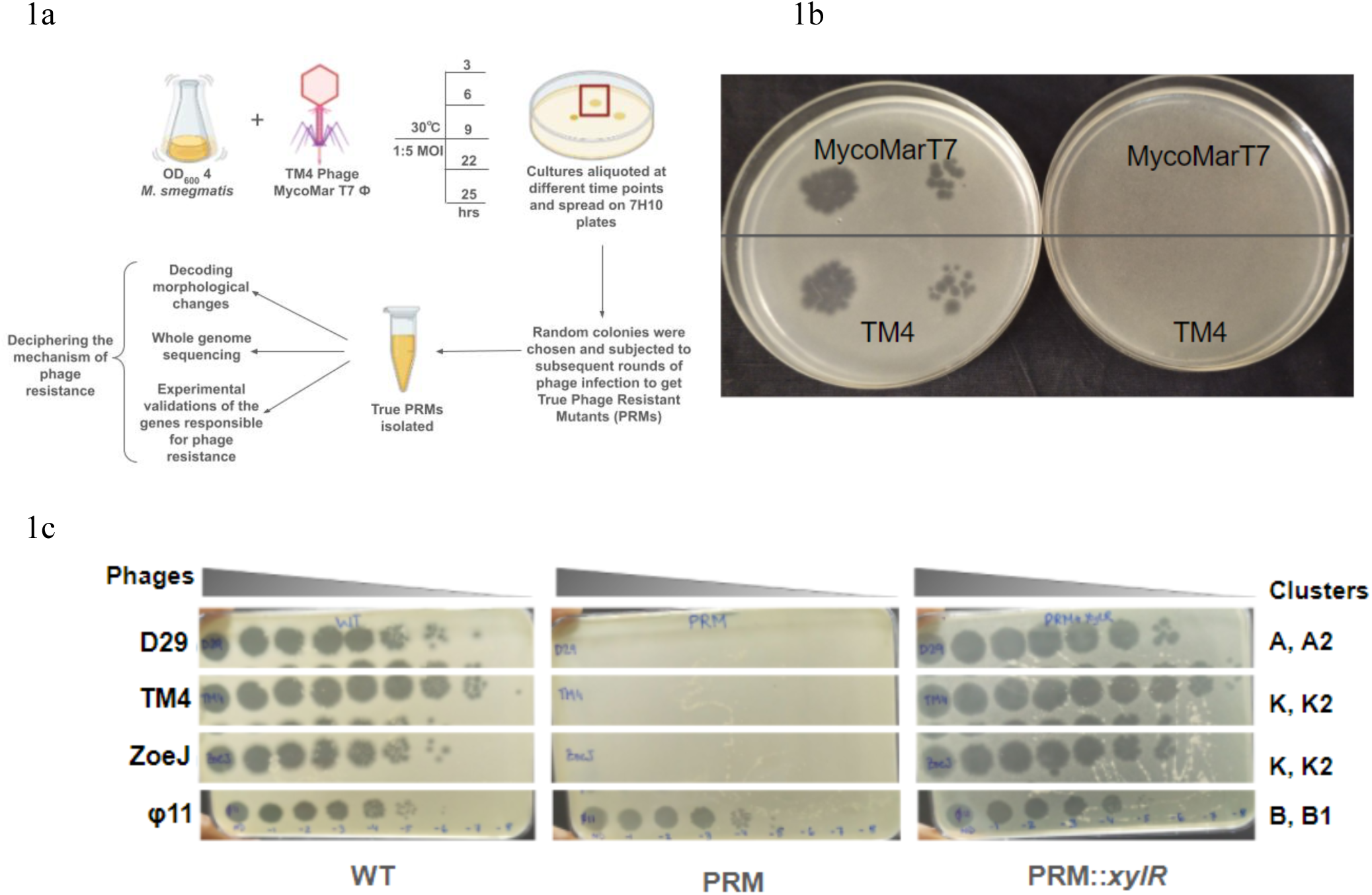
*M. smegmatis* mutants resistant to TM4 Mycobacteriophage. **(a)** Schematic representation of generating spontaneous phage-resistant mutants in *M. smegmatis*. **(b)** Plaque assay showing the sensitivity of wild-type (WT) *M. smegmatis* and the phage-resistant mutant (PRM) to MycoMar T7 phage. Serial dilutions of the phage lysate were spotted onto bacterial lawns. Clear plaques were observed on WT lawns, whereas no cell lysis was detected on PRM, indicating resistance to MycoMar T7 phage. **(c)** Phage susceptibility of WT *M. smegmatis mc²155*, PRM, or PRM::*xylR* strains towards different Mycophages. Tenfold serial dilutions of phage lysates (D29, TM4, ZoeJΔ43–45 [ZoeJ], and PDRPxv [Phage11]) were spotted on solid media with bacterial lawns. Plaque assays were performed at least three independent times with consistent results.

Given that MycoMarT7 is derived from TM4, we next evaluated whether the PRM exhibited similar resistance to the parental TM4 phage. Plaque assays revealed that PRM was equally resistant to TM4, showing no zones of clearance, consistent with its MycoMarT7 resistance profile (Figure 1b). Based on these results, TM4 was used for all subsequent experiments. Interestingly, the PRM displayed a broader resistance beyond TM4. To test for cross-resistance to various phages, we challenged the mutant with a panel of genetically diverse mycobacteriophages, such as D29 (cluster A2), ZoeJΔ43-45 (cluster K2)^40^, PDRPxv (cluster B1)^41^. These phages represent multiple unrelated genetic clusters, each utilizing distinct infection strategies and host determinants. Remarkably, PRM exhibited complete resistance to most of the tested phages, expect for Phage 11, as evidenced by the absence of plaque formation or lytic clearing in spot assays (Figure 1c). This phenotype indicates that the resistance mechanism is not only cluster specific but phage specific. Therefore, our observation suggests that mutation conferring resistance in PRM likely target a fundamental aspect of the phage infection cycle rather than a receptor or function unique to a single phage. This broad resistance profile makes PRM a valuable model for dissecting the genetic and physiological determinants of mycobacterial phage susceptibility.

### Phage resistant mutant displays altered surface phenotype

Beyond its resistance to mycobacteriophage, PRM displayed several phenotypic traits that clearly distinguished it from the parent *M. smegmatis* mc²155 strain that includes smaller, smooth, and glossy colonies, compared to the larger, wrinkled and rough morphologies associated with *M. smegmatis* (Figure 2a). This altered colony architecture suggested changes in cell surface structure and/or extracellular matrix production. To understand whether the smaller colony size was associated to a growth defect, we studied bacterial growth kinetics in liquid culture. PRM exhibited a slower growth rate compared to WT and consistent with smaller colonies on agar (Figure S1a).

**Figure 2:**
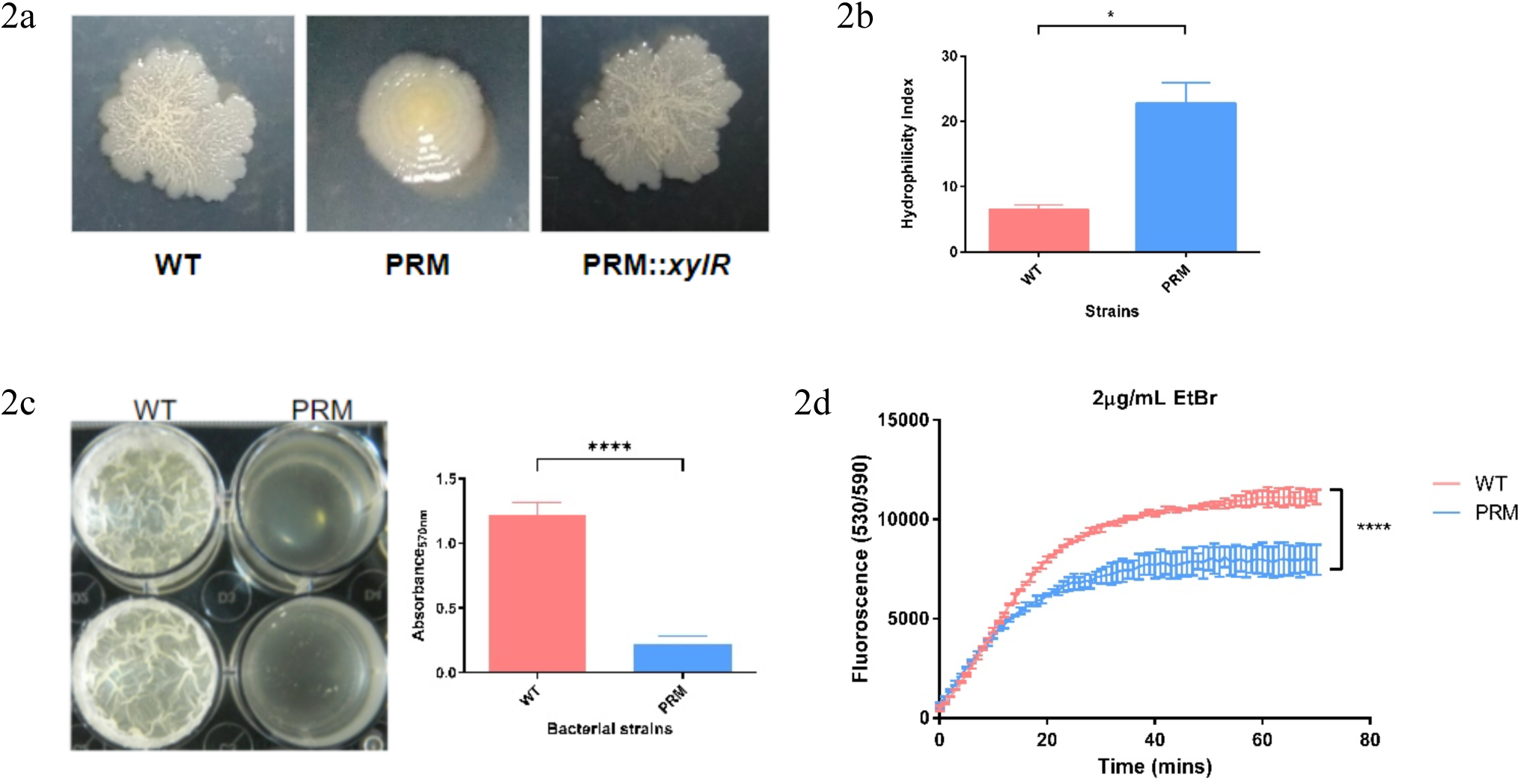
The spontaneous mutant has altered phenotype. **(a)** Colony morphology of the WT *M.smegmatis*, PRM and PRM::*xylR* spotted on 7H10 solid medium **(b)** Hydrophilicity assay between WT and PRM **(c)** Biofilm formation in M63 medium and quantitation by staining with crystal violet **(d)** Differential EtBr uptake between WT and PRM cells. All the tests were performed in three biological replicates, and error bars indicate the standard deviations. Statistical analysis was carried out using multiple Student’s *t*-tests. The *P*-values of the results (<0.05, <0.0001) are indicated by asterisk (*, ****).

Given the morphological differences, we next examined the physiological properties of the bacterial cell surface. A hexadecane partitioning assay revealed that PRM cells were more hydrophilic than WT, as indicated by a reduced partitioning into the organic phase (Figure 2b) consistent with a ruffled cellular morphology (Figure S1b). Pellicular biofilm, a surface-associated phenotype strongly influenced by cell envelope composition, was evaluated under static growth conditions. The PRM strain demonstrated a defect in biofilm formation at the air-medium interface as compared to WT and with a significant reduction in biofilm mass (Figure 2c), indicating a compromised ability to form a robust biofilm. We further investigated whether the envelope-related phenotypic changes were accompanied by alterations in membrane permeability. By using the ethidium bromide (EtBr) uptake assay, we observed that PRM cells accumulated the dye much slower than wild type cells, suggesting reduced membrane permeability (Figure 2d). Reduced permeability limits the entry of hydrophilic molecules, including antibiotics and possibly impede the passage of phage DNA, through the inner layers of the cell envelope.

To explore whether altered membrane permeability leads to changes in antimicrobial susceptibility, we determined the susceptibility against several anti-TB drugs. The MICs for most antibiotics, including streptomycin, rifampicin, moxifloxacin, norfloxacin, and ethambutol, remained unchanged. However, a modest (2-fold) increase in resistance to isoniazid was observed in the mutant. In contrast, the mutant exhibited increased susceptibility, reflected by reduced MICs (2-fold), to kanamycin and D-cycloserine (Figure S1c). Consistent with this, cell viability assays showed a 10 fold higher killing with kanamycin (Figure S1d). These findings suggest selective changes in susceptibility between wild type and PRM, though the underlying mechanisms remain unclear and warrant further investigation. This also indicates that uptake of specific antibiotics are affected by the altered membrane permeability in PRM and therefore emergence of phage resistance could synergize or antagonize with antibiotics. Taken together, these findings demonstrate that PRM displays an altered phenotypes including surface morphology, slower growth, increased hydrophilicity, impaired biofilm formation, and decreased membrane permeability, that are likely interconnected. This also highlight the complex interplay between phage defense mechanisms and cell envelope dependent features in *M. smegmatis*.

### Mutation in *xylR* contributes to multiphage resistance in *M. smegmatis*

To identify the genetic basis of the altered phenotypes in PRM, its genome was sequenced and comparison with the reference strain revealed a 12 nucleotides in-frame deletion within the *xylR* gene. *xylR,* is a xylose-responsive transcriptional repressor in *M. smegmatis*, is known for its role in regulating the operon for xylose catabolism. Along with regulating xylose metabolism in the cells, recent studies have found that *xylR* also regulates the expression of around 1,500 genes, suggesting its broader influence on cellular physiology, including lipid metabolism pathways, stress response, and cell envelope structure among others^25^. Given the scope of *xylR*-mediated transcriptional regulation, we hypothesized that a 12 nucleotide deletion in *xylR* may have caused its loss of function, leading to phage resistance and the altered phenotype observed in PRM. To test this hypothesis, a wild type copy of *xylR* was cloned under a constitutive promoter and overexpressed in PRM (referred to as PRM::*xylR*). Upon overexpression of XylR, the complemented strain displayed a complete reversal of the mutant phenotype. The smooth and shiny colony morphology of PRM reverted to the characteristic rough and wrinkled appearance, indicating restoration of the native surface architecture (Figure 2a). While PRM remained resistant to TM4 and other phages, PRM::*xylR* strain showed appearance of plaques similar to that of the with type strain, indicating restored sensitivity to TM4 infection (Figure 1c), and demonstrating that loss of *xylR* activity in PRM was responsible for the resistance.

To further assess the dynamics of phage infection across these strains, we performed time-course kill curve assays. Cultures of wild type mc²155, PRM, and PRM::*xylR* were infected with TM4 at an MOI of 1:10 in liquid medium, and viable colony-forming units (CFUs) were measured at regular intervals post-infection. Both WT and PRM::*xylR* exhibited a sharp decline in viability over time, indicative of productive phage infection and host lysis. In contrast, PRM maintained high CFU counts throughout the course of infection, consistent with effective resistance to TM4-mediated killing (Figure 3a). These findings were further supported by spot plating assays on solid media containing TM4 phage. Serial dilutions of each strain were spotted on plates with excess phages. As expected, PRM was able to grow robustly even in the presence of phage, whereas neither WT nor PRM::*xylR* could form colonies under these conditions (Figure 3b). Taken together, these results confirm the deletion in *xylR* as the cause for conferring resistance to TM4 and an altered colony morphology observed in PRM. Our findings implicate *xylR* as a key node in the genetic network linking transcriptional regulation, cell surface remodeling, and phage-host interactions in *M. smegmatis*.

**Figure 3:**
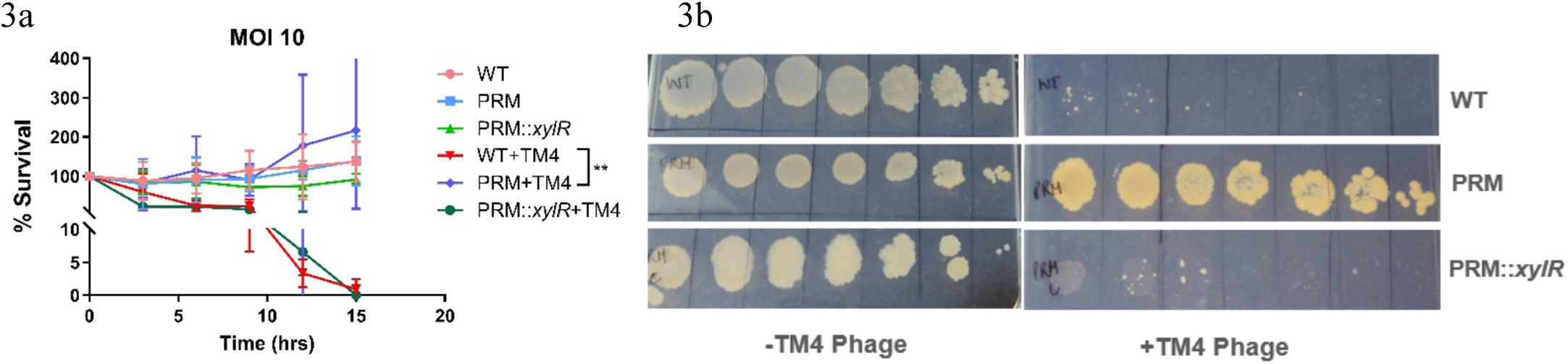
Mutation in *xylR* imparts phage resistance in *M. smegmatis*. **(a)** Bacterial viability in presence and absence of TM4 phages at MOI 1:10. The infection was performed in liquid culture for 15 hrs. Samples were collected every 3hrs and CFU was enumerated. All the tests were performed in three biological replicates, and error bars indicate the standard deviations. Statistical analysis was carried out using multiple Student’s *t*-tests. The *P*-values of the results (<0.01) are indicated by asterisk (**). **(b)** Bacterial survival on phage lawns. TM4 phage was incorporated into top agar to form a phage lawn, on which tenfold serial dilutions of WT, PRM, and PRM::*xylR* cells were spotted to assess their ability to survive and grow in the presence of phage. All the tests were performed in three biological replicates, and error bars indicate the standard deviations.

### Structural analysis suggests impaired DNA binding ability in the mutant XylR

To understand the impact of in frame deletion (^162^LVHG^165^) in XylR on its function it is crucial to understand the structural architecture of the wild type protein and compare it with its variants. Pfam domain analysis of the XylR protein revealed the presence a C-terminal ROK domain spanning from residue 105-379 with potential involvement in sugar metabolism and transcriptional repression. The characteristic features of ROK family proteins include the presence of a DNA binding domain, ATP binding motif and Zinc-binding cysteine motif^42^. N-terminal residues 25-73 in XylR showed the presence of a conserved helix-turn-helix motif while the rest of the N-terminal domain contained predominantly helical conformation supporting its role in DNA binding. The molecular structure of the wild-type XylR protein was modeled through homology modelling using *E. coli* Mlc protein (PDB ID-1z05.1.A) as a template structure. The modelled structure shows XylR protein exists in the homo-dimeric form that is consistent with the architecture of the ROK family transcription regulators (Figure 4a, S2a). The dimerization of the XylR protein was mediated through intrachain H-Bonds formed between the residues M320-G249 (2.9 Å), and N319-H253 (3.3 Å) at interchain interaction region.

**Figure 4:**
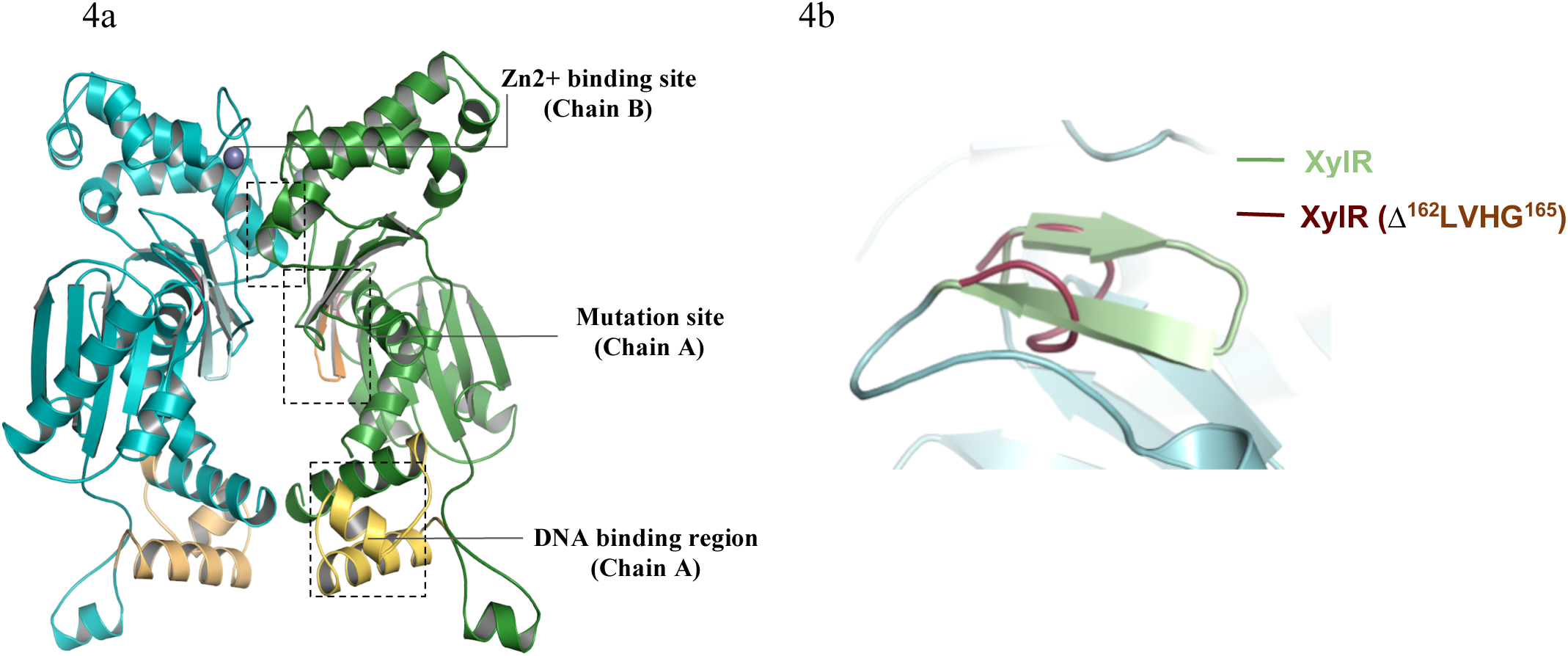
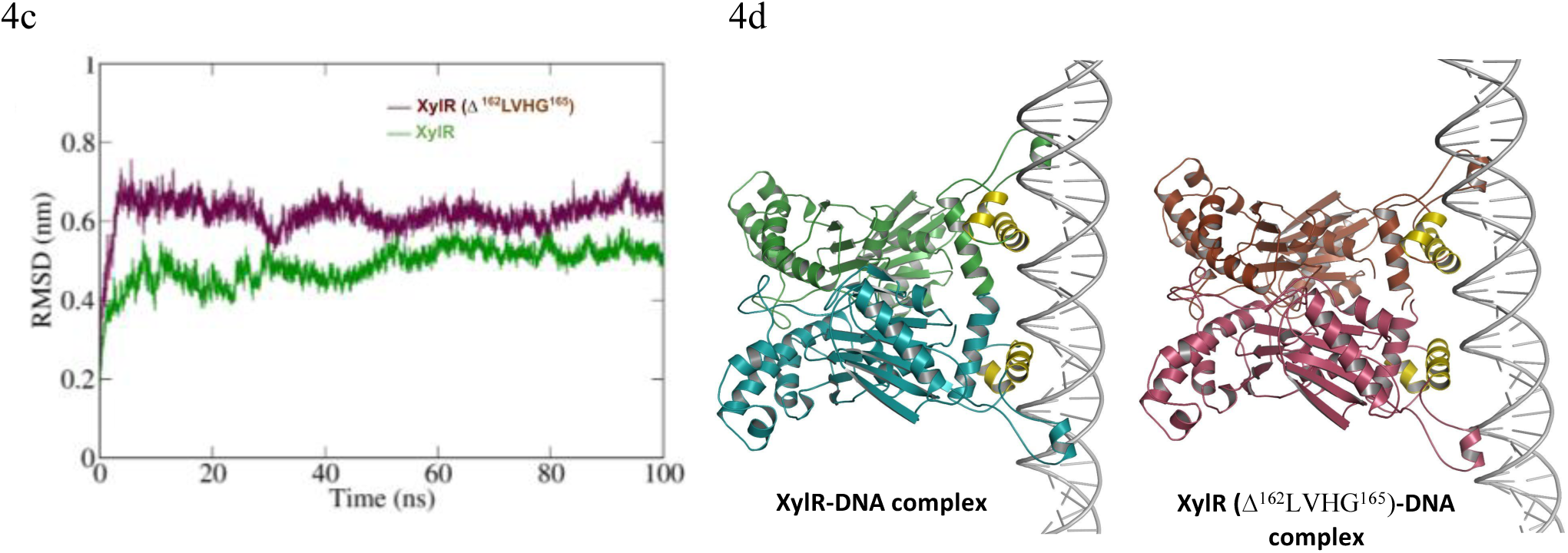
Structural characterization and the molecular dynamics simulation analysis of WT and Δ^162^LVHG^165^ XylR protein. **(a)** The molecular structure of the XylR homodimer (Chain A shown in cyan, Chain B in green) indicating DNA-binding region, site of mutation site, and Zn²⁺ binding site. **(b)** Loss of secondary structural element in the mutant protein compared to the wild type. **(c)** The backbone RMSD of the XylR and Δ^162^LVHG^165^ XylR proteins over 100ns simulation time, indicating structural stability during MD simulation. **(d)** HADDOCK-based protein-DNA docking analysis of XylR (left panel) and Δ^162^LVHG^165^ XylR (Right panel).

The observed ^162^LVHG^165^ deletion was induced in XylR and the mutant structure was subjected for local energy minimization to counter the steric strain. Upon structural superimposition, it was observed that the antiparallel β-sheets at site of deletion turned into a flexible loop region in the mutant protein (Figure 4b). This disruption of secondary structure at the hinge region suggests potential impairment in domain communication and conformational stability of the mutant XylR protein. 100 ns MD simulation was performed to verify the impact of the secondary structure disruption on the overall structure of XylR. Furthermore, through the course of simulation, the disrupted β-sheets at the site of mutation in XylR (Δ^162^LVHG^165^) did not regain the native conformation (Figure S2b). Root Mean Square Deviation analysis showed that the wild-type protein attains the convergence with average deviation of approximately 0.5nm, whereas the mutant protein exhibited increased deviation of 0.6nm (Figure 4c, S2c). Additionally, a PCA analysis of the MD simulations revealed changes in the collective motion pattern between wild type and mutant protein. (Figure S2d). To further our understanding on the effect of these structural alterations on the DNA binding ability of XylR we performed flexible docking of the dimeric XylR on the promoter DNA sequence of the xylose operon. Expectedly, the interaction of the third helix (H3) of the HTH motif with the major grove of the promoter DNA was found to be impaired in case of the mutant (Figure 4d, S2e). Mapping of the atomic interactions between DNA and the homodimeric repressor showed a marked reduction in H-bonds in XylR (Δ^162^LVHG^165^)-DNA complex. In case of XylR, both chains established multiple stable interactions with the promoter DNA, supporting efficient binding while XylR (Δ^162^LVHG^165^) exhibited a marked asymmetry in binding, with chain B completely lacking detectable H-bond interactions with the DNA (Figure S2e). This loss of stability of the repressor–DNA complex, results in impaired DNA-binding affinity and consequent loss of transcriptional regulation.

### Phage attachment and intracellular replication are not affected in PRM

Given the differences in cell surface characteristics between the wild-type and PRM, we sought to determine whether these modifications influenced TM4 attachment to *M. smegmatis*. To this end, we performed a quantitative adsorption assay at a MOI of 1:0.01. As a negative control for adsorption, WT cells were incubated with the surfactant Tween 80, known to inhibit phage binding to bacterial cell surfaces. In WT+Tween 80 control, the percentage of unadsorbed phages remained essentially constant over the assay period, confirming the inhibitory effect of Tween 80 on TM4 binding. In contrast, all three experimental strains—WT, PRM, and PRM::*xylR*—showed a time-dependent decline in the proportion of unadsorbed phages (Figure 5a) suggesting that phage binding is not affected in PRM. Notably, there was a decrease in phage titre post 20 minutes, with each strain demonstrating progressive phage adsorption. Although the magnitude of reduction varied slightly among strains, TM4 binding was not impaired in PRM or in the complemented strain compared to the WT, suggesting that the deletion in *xylR* (^162^LVHG^165^) in PRM does not hinder initial phage attachment. To validate these findings, we also performed flow cytometry analysis to study TM4 binding at the single-cell level. TM4 phages were stained with Sytox green dye, allowing detection of phage-bound bacteria via FITC fluorescence. Binding was assessed by measuring an increase in side-scatter, and Sytox green (FITC) signal. Increases in side scatter reflects changes in cell surface complexity due to phage attachment, while FITC intensity corresponds to the fluorescence signal from bound Sytox green–labelled phages. Similar to phage adsorption assay, we observed a time-dependent increase in both side scatter and FITC values for all the strains (Figure 5b, S3a) indicating binding of phage particles to the bacteria. Importantly, the extent of side scatter and FITC increase in the PRM closely resembled with WT, further supporting that adsorption of TM4 phage to the mycobacterial cell surface is unaffected by the loss of *xylR* function.

**Figure 5:**
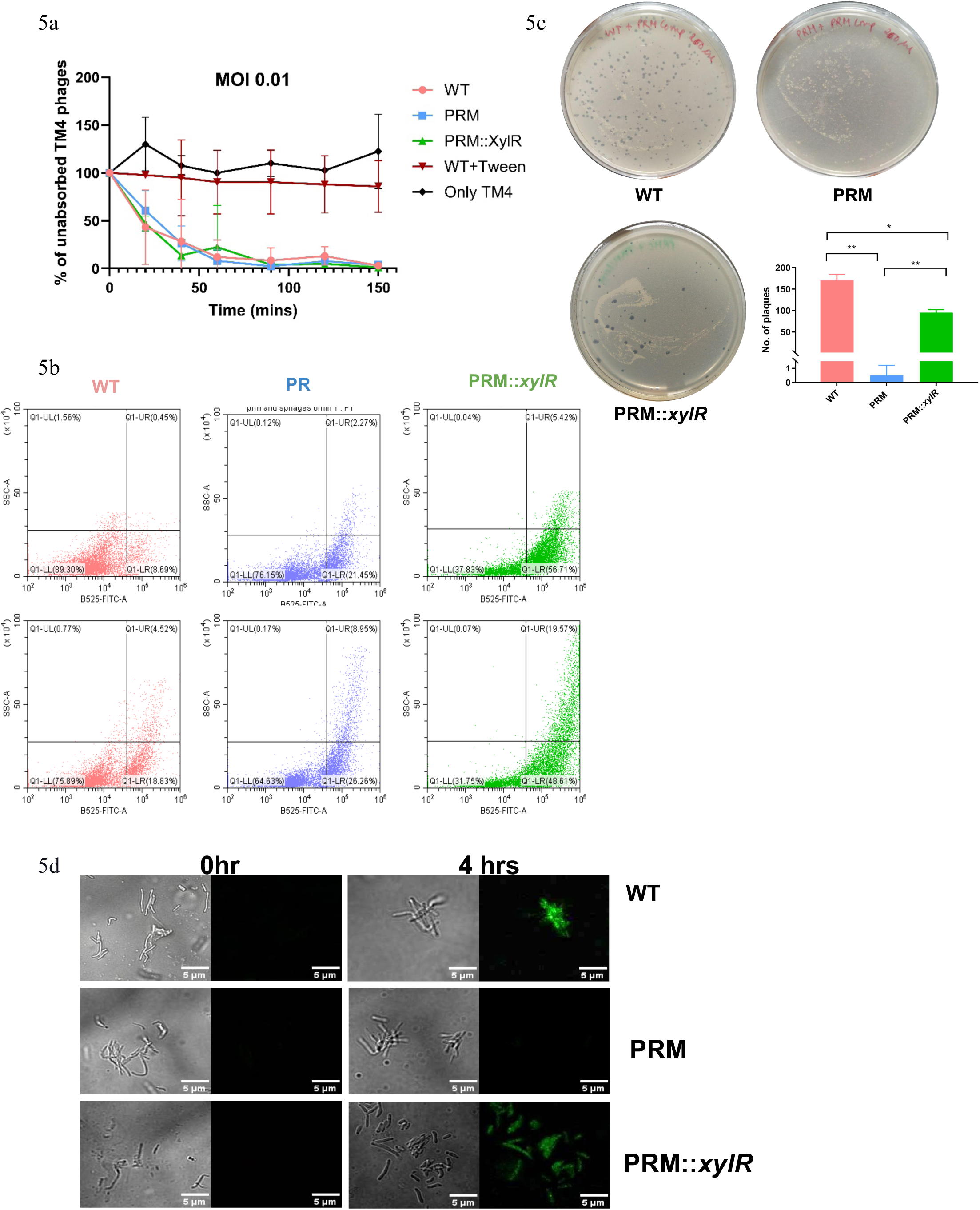
TM4 Phage attachment and its intracellular life cycle in *M. smegmatis*. **(a)** Comparison for adsorption of phage TM4 to different mycobacterial strains, including WT, PRM, or PRM::*xylR*. All the tests were performed in three biological replicates, and error bars indicate the standard deviations. **(b)** Flow cytometry analysis showing percentage of bacilli bound by the TM4 phage labelled with Sytox green, relative to the study population. This assay was conducted twice. **(c)** DNA electroporation assays for determining if the phage genomic DNA can complete the subsequent life cycle after entry to the mycobacterial cell. The extracted TM4 genome was electroporated into the PRM strain. After 3 h incubation, this sample was mixed with the WT, PRM, or PRM::*xylR* strains, and the number of plaques formed were assessed. All the tests were performed in three biological replicates, and error bars indicate the standard deviations. Statistical analysis was carried out using multiple Student’s *t*-tests. The *P*-values of the results (<0.05, <0.01) are indicated by asterisk (*, **). **(d)** TM4 Phage DNA Injection Post-Adsorption to *M. smegmatis* cells. Microscopy of WT, PRM, or PRM::*xylR* strains infected with fluorophage φ^2^GFP10. For PRM no fluorescence is observed even after 4hrs, indicating lack of successful phage infection. This experiment was performed 3 independent times.

Since phage binding to PRM is not the cause of phage resistance, we next evaluated whether post-injection phage life-cycle steps are impaired in PRM. To check this, we performed electroporation assays to examine if the TM4 phage genomic DNA remain stable upon entering the mycobacterial cell and was able to complete full life cycle and lyse the host cells. When the genomic DNA of TM4 phage (Figure S3b) was electroporated into the PRM strain and plated after mixing with the three different strains, wild-type, PRM and PRM::*xylR*. Plaques were observed in the case of wild-type and PRM::*xylR* strain. In contrast, no plaque were formed on the plate mixed with PRM strain (Figure 5c), indicating that the resistance phenotype in PRM is not due to defects in post-injection life-cycle events such as intracellular phage DNA replication or virion maturation, but rather to a defect at an earlier stage. These results confirm that, once TM4 genomic DNA enters mycobacterial cells, the subsequent stages of the phage life cycle including replication, assembly, and release proceeds normally, generating infectious phage particles capable of forming plaques on susceptible hosts. Taken together with adsorption assay, our findings suggest that while TM4 can attach to the PRM cell surface, the delivery of phage genomic DNA into the host cytoplasm is impaired, preventing a productive lytic infection. It is also possible that phage attachment to its host cell is a two-step sequential procedure where first a non-specific attachment of the phage takes place followed by a correct receptor binding and successful release of the phage genome. Improper receptor binding may result in unsuccessful genome delivery despite showing phage-Mycobacteria adsorption.

### TM4 Phage DNA injection is dependent on *xylR*

Since PRM strain was not deficient in binding to TM4 phages or restrict the replication and assembly of TM4 virions, we explored if the step of phage DNA injection is inhibited in the *xylR* mutant. To directly assess whether TM4 phage DNA successfully enters the host cytoplasm in the PRM, we employed the fluorophage φ^2^GFP10^43^. This engineered derivative of TM4 encodes a green fluorescent protein that is expressed during viral particle packaging and only becomes detectable at the late phase of the life cycle. Thus, GFP expression serves as a reliable reporter of successful phage genome entry, replication, and assembly within the bacterial cell^43^. Previous work has reported that the TM4 lytic cycle in *M. smegmatis* is completed within approximately 4 hours post-infection. Therefore, WT and PRM cells were infected with φ^2^GFP10 at MOI of 0.01 and imaged at 0, 3 and 4 hr post-infection. Expectedly, uninfected WT cells exhibited no green fluorescence at time 0, while at 3 hours, a subset of WT cells (data not shown), and by 4 hours, the majority of cells displayed a strong GFP signal, indicating that productive infection in a substantial fraction of the population (Figure 5d). In contrast, PRM cells remained completely non-fluorescent at all-time points. The absence of GFP expression in PRM indicates that the phage genome did not successfully enter the cytoplasm to initiate its transcription and replication. This blockade in the infection process occurs downstream of adsorption, consistent with a failure at the DNA injection stage and confirms that an intact XylR function is essential for facilitating TM4 phage DNA delivery into the host cell, and that its disruption results in complete resistance to infection despite normal phage binding.

### PRM has a higher basal metabolic rate as compared to that of wild type *M. smegmatis*

Given that *xylR* functions as a global transcriptional repressor, we hypothesized that disruption of the DNA binding ability due to the deletion of ^162^LVHG^165^ in PRM may lead to the de-repression of downstream target genes which contribute to the resistant phenotype. To test this, we performed comparative transcriptomic analysis of the mutant. We first assessed the overlap between biological replicates of WT and PRM samples to ensure dataset reliability. Analysis showed that PRM exhibited a broad transcriptional remodeling, consistent with the disruption of a global transcriptional regulator. We applied a log_2_ fold change cutoff of ±1.5 among the 6017 differentially expressed genes (Figure 6a, S4, TableS1). Functional categorization of all the differentially expressed genes revealed a prominent upregulation of genes associated with ABC transporters, membrane transport systems, conserved hypothetical proteins, and cell wall–related processes in the PRM. Notably, a substantial proportion of upregulated genes were involved in intermediary metabolism and respiration. These findings indicate that disruption of *xylR* drives extensive metabolic rewiring, leading to broad alterations in cellular physiology and overall bacterial function (Figure 6b, TableS1).

**Figure 6:**
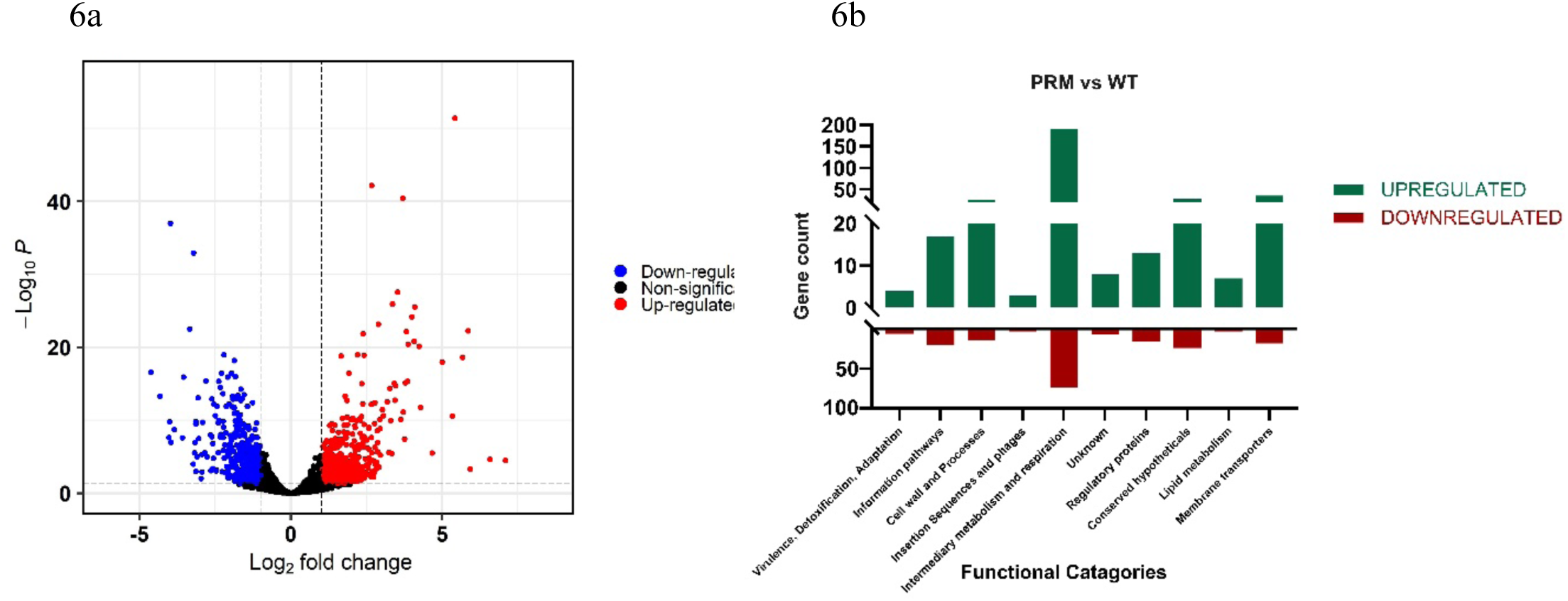
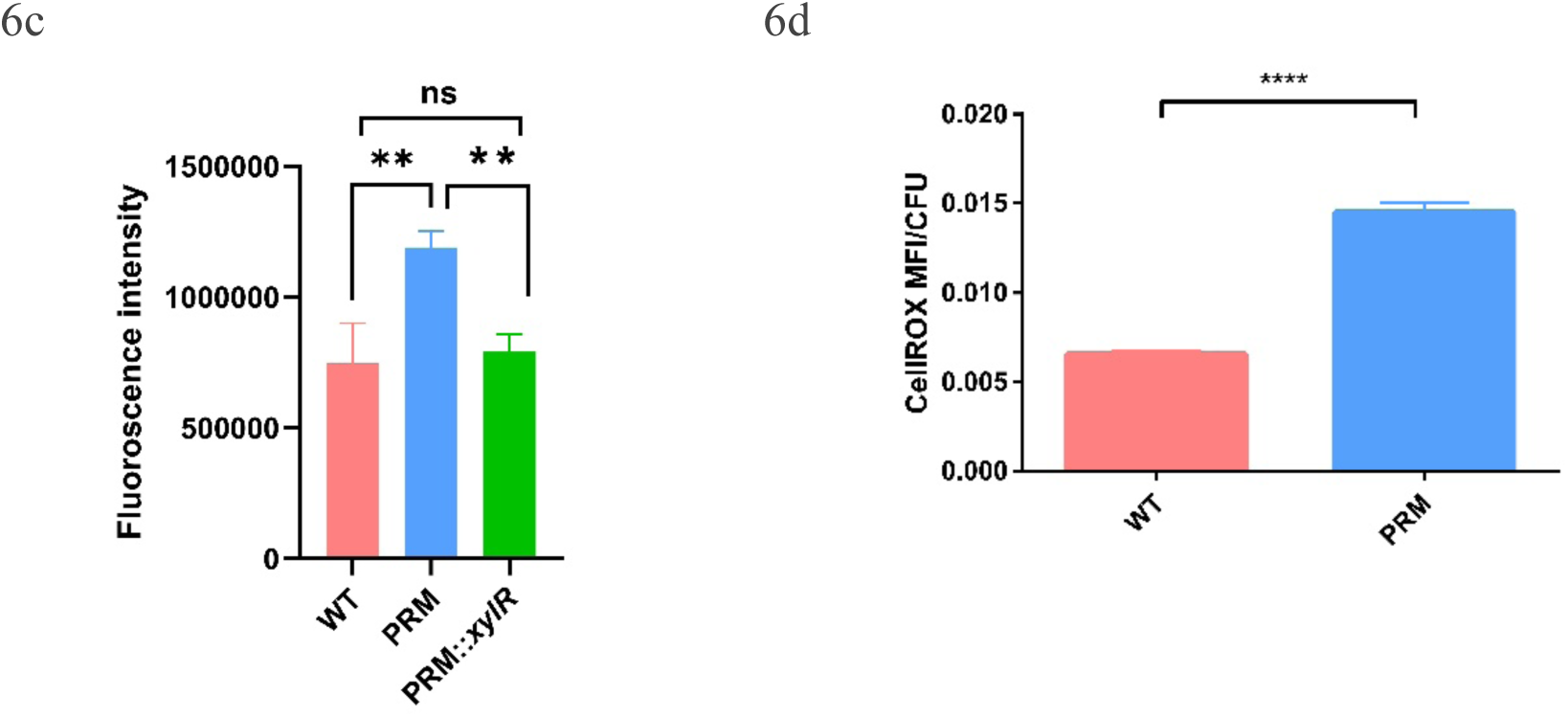
Transcriptomics and metabolic analysis of PRM and wild type *M. smegmatis* mc^2^155. (a) Volcano plot shows gene expression differences between WT and PRM strains through transcriptomic analysis with significant down-regulated (blue) and up-regulated (red) genes. (b) Functional categories of the upregulated and downregulated genes. Fold change of ≥ 1.5 was chosen for selecting the differentially expressed genes. (c) Resazurin reduction assay showing bacterial respiration rate. The fluorescent intensity shows the rate of conversion of Resazurin to resorufin. All the tests were performed in three biological replicates, and error bars indicate the standard deviations. (d) Mean fluorescence intensities of CellROX deep red dye with respect to CFU between WT and PRM strains. Statistical analysis was carried out using multiple Student’s *t*-tests. The *P*-values of the results (<0.01, <0.0001) are indicated by asterisk (**, ****), and ns- not significant difference.

These findings also suggested that PRM exhibits a higher metabolic activity as compared to WT. To validate this observation, we performed a resazurin reduction assay. Resazurin is a blue, non-fluorescent compound that is reduced to a pink, fluorescent resorufin in metabolically active cells. Increased fluorescence intensity reflects enhanced respiratory activity. PRM displayed significantly higher fluorescence compared to WT and the complemented strain, indicating increased metabolic activity (Figure 6c). As elevated metabolism is often associated with an increase in the production of reactive oxygen species (ROS), we next measured basal ROS levels using CellROX Red dye. PRM cells exhibited significantly higher ROS levels per cell relative to WT (Figure 6d), thus collectively suggesting a higher metabolic rate in PRM. However, these data did not clarify how the observed transcriptional changes confer phage resistance. To further elucidate the mechanism, we performed transcriptomic analysis of TM4-infected bacterial strains.

### XylR directly regulates expression of the LOS synthesis gene island

Cells infected with TM4 were harvested at 140 minutes before the completion of the lytic cycle (180 min), and total RNA was subjected to sequencing. CFU were measured to confirm absence of cell death within this time (Figure S5a, TableS1). Comparative gene analysis showed a high overlap in the global gene expression distribution between the two strains (Figure S5b, TableS1), supporting the reliability of the dataset. Importantly, no significant differences were observed in the induction of genes implicated in canonical anti-phage defense or in stages of the TM4 infection cycle such as adsorption, phage DNA replication, or virion assembly. Consistent with the phenotypic assays showing that these stages of the TM4 infection cycle remain intact in the PRM. Expectedly, in case of the wild type strain, we observed an increase in the transcript count from the TM4 genes, which were dominated by the presence of late phase genes of the phage replication cycle. To our surprise, a rise in the transcript count of the phage genes were also recorded in case of PRM, indicating the inability of PRM to fully resist the infection by TM4 phage at very high titre (MOI of 1:10). We then tracked the expression level of the bacterial genes that changed during the course of infection, phage replication and packaging in the wild type before the onset of lysis. Thereafter, we compared the change in the expression pattern with those in the mutant. Notably, we found a good correlation between the gene expression between the two strains (Figure 7a, 7b, TableS1) confirming the occurrence of successful infection in PRM following the same sequence of replication when challenged with a very high titre of TM4 phages. Our data suggests that the resistance is not mediated by canonical antiviral defense systems but rather might arise from *xylR*-dependent transcriptional reprogramming that promotes the formation of a modified, polar lipid-rich cell envelope also responsible for increased surface hydrophilicity in PRM. A striking feature of the transcriptomic data was the significant upregulation of 6 genes within the cluster comprising MSMEG_4727–MSMEG_4737, annotated as the lipooligosaccharide (LOS) biosynthesis gene island (Figure 7c), suggesting an increase in LOS production in PRM. Enhanced LOS production modifies the physico-chemical properties of the mycobacterial cell surface, potentially altering membrane organization at the phage–host interface. Although the phages efficiently bind to the mutant cells, we hypothesised that if PRM has accumulated excessive amounts of LOS then it may impede the membrane penetration steps required for successful phage DNA injection. These data are supported by an absence of phage genome replication and virion assembly in PRM despite normal adsorption, thereby localizing the defect to an early post-adsorption stage of infection.

**Figure 7:**
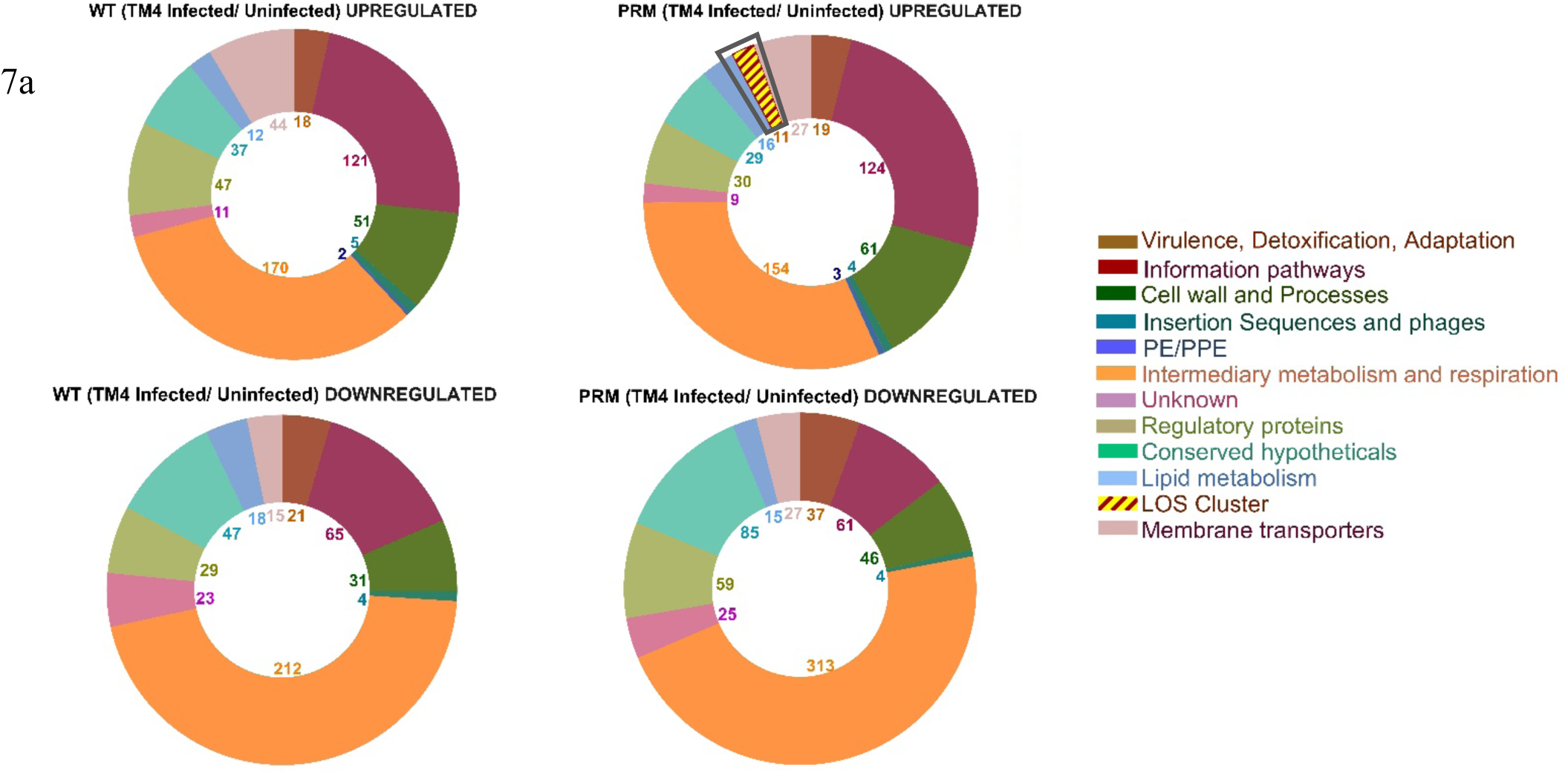
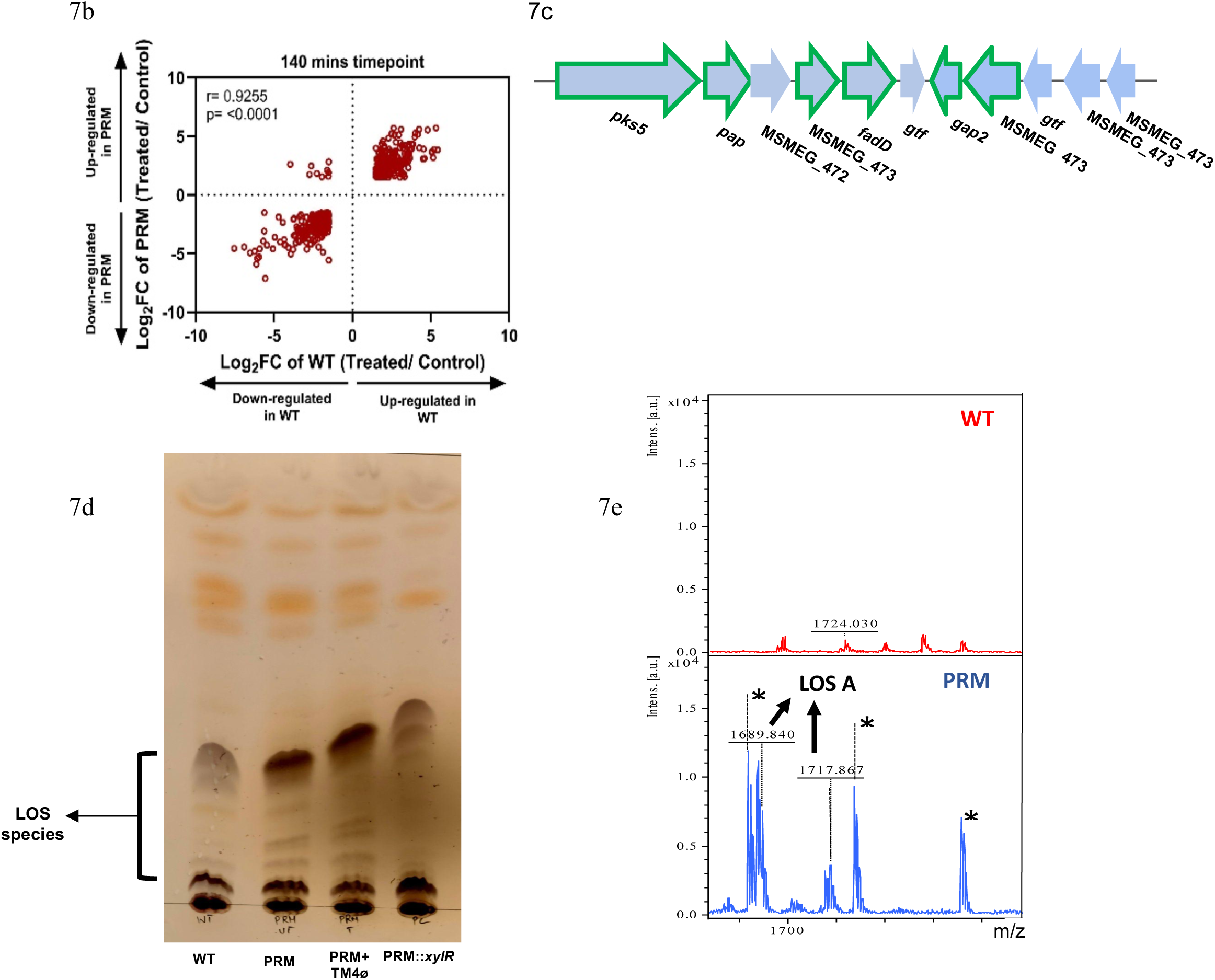
Regulation of LOS synthesis gene island by *xylR*. **(a)** Functional categories of the genes (TM4 Phage Infected/Uninfected) upregulated in WT (Top left), PRM (Top right), and downregulated in WT (Bottom left), PRM (Bottom right) (FC ≥1.5). Genes representing to the LOS gene cluster are highlighted in yellow-maroon pattern. **(b)** Scatter plot of the Pearson correlation analysis of gene expression in PRM and WT upon TM4 phage infection. Each dot represents a single gene with its log_2_ fold difference for WT (x-axis) and PRM (y-axis). **(c)** Pictorial representation of the LOS gene cluster. Out of the 11 genes, the ones highlighted in green are the ones showing significant upregulation in the transcriptomic data. **(d)** Thin layer chromatography analysis of mycobacterial membrane lipids. Lipids were extracted from exponentially grown cultures of *M. smegmatis* strains, resolved on the solvent (CHCl_3_-CH_3_OH, 90:10) and visualized by spraying with alpha napthol, followed by charring. Positions of the LOS are indicated with an arrow. **(e)** MALDI-TOF MS analysis of the lipid extracts of the WT, and PRM strains. [M + Na]^+^ adducts of LOS-A are labelled. The repeat peaks differing by 44 Da (marked with *) originates from a contaminating PEG species from the plasticware used for MALDI-TOF MS.

To determine whether disruption of *xylR* affects the glycolipid composition, total lipids from WT, PRM, PRM infected with TM4 phage, and PRM::*xylR* were extracted and analysed with a LOS specific solvent system using thin layer chromatography (TLC) (Figure 7d). Few lipid species were markedly enriched in both the PRM conditions compared to WT (lanes 1, 2 and 3). Based on earlier reports, this band likely correspond to lipooligosaccharides (LOS), specifically the dipyruvylated LOS-A forms with varying acyl chains^37^. Additionally, two minor bands with lower mobility were also enriched in the PRM and are consistent with the polar monopyruvylated LOS-B1 and LOS-B2 species^31^. Notably, these LOS-associated bands were mostly absent in the WT and PRM::*xylR* strains (lane 4). Quantitative analysis of the TLC further revealed significant upregulation of LOS species in TM4-infected PRM relative to untreated PRM, with minimal levels observed in WT (Figure S5c). To confirm the identity of the bands enriched in PRM, the glycolipids were subjected to MALDI-TOF mass spectrometric analysis following purification by preparative TLC. PRM lipid extracts, MALDI-TOF analysis revealed three major clusters of [M + Na]⁺ adduct ions: with m/z 1159.64-1229.71, corresponding to type II A glycopeptidolipids (GPLs), a second cluster at m/z 1329.68–1371.66 corresponding to type III A GPLs^44^, and a third cluster at m/z 1689.84 – 1717.86, which matched with the calculated masses of [M + Na]⁺ adduct ions of LOS-A species^45,46^ (Figure 7e, S5d). This LOS-associated cluster was absent or barely detectable in WT total lipid extracts but was clearly enriched in the PRM. Taken together, these results suggests that the loss of function of XylR transcriptionally activates LOS biosynthesis and promotes increased accumulation of LOS on the outer cell envelope, likely increasing surface hydrophilicity or complexity and thereby blocking TM4 phage infection at an early post-adsorption step, most plausibly during phage DNA injection.

### Chemical treatment of PRM reverses resistance to TM4 phage

To determine whether the phage-resistant phenotype could be reversed by externally perturbing the modified cell surface, we sought conditions that would help in the removal of this excess outer lipid layer predominated by LOS, while preserving cell viability. Rather than employing harsh chemical treatments, we tested whether exposure to polyols or osmolytes such as glycerol, sorbitol, or PEG could facilitate peeling of the surface-associated material. Remarkably, washing PRM cells with these osmolytes restored phage susceptibility, as evidenced by the reappearance of plaques (Figure 8a). This restoration of phage sensitivity indicates that the resistance phenotype is not due to an irreversible structural alteration in the core cell wall, but rather due to the composition of the surface-exposed and loosely associated lipid layer that can be modulated under specific conditions. To directly assess the involvement of LOS, we analysed the extracted non-covalently attached lipids by TLC after treatment with polyols. The treated PRM samples showed a significant decrease in LOS-specific bands compared to that of untreated PRM cells (Figure 8b), suggesting a partial removal of the surface-associated LOS. These findings strongly support a functional role for LOS in mediating phage resistance. Taken together, these results demonstrate the accumulation of excess LOS on the cell surface that acts as a hindrance for successful phage infection. *xylR*-dependent upregulation of LOS biosynthesis produces a protective envelope that allows phage adsorption but hampers subsequent DNA release into bacteria. Importantly, removal of excess LOS restores phage susceptibility, substantiating our conclusion that LOS enrichment underlies resistance to TM4 infection. Together, our findings provide a novel mechanism of mycobacterial resistance to phage, which could aid in better understanding of phage-bacterial interactions and development of potential therapeutic strategies.

**Figure 8:**
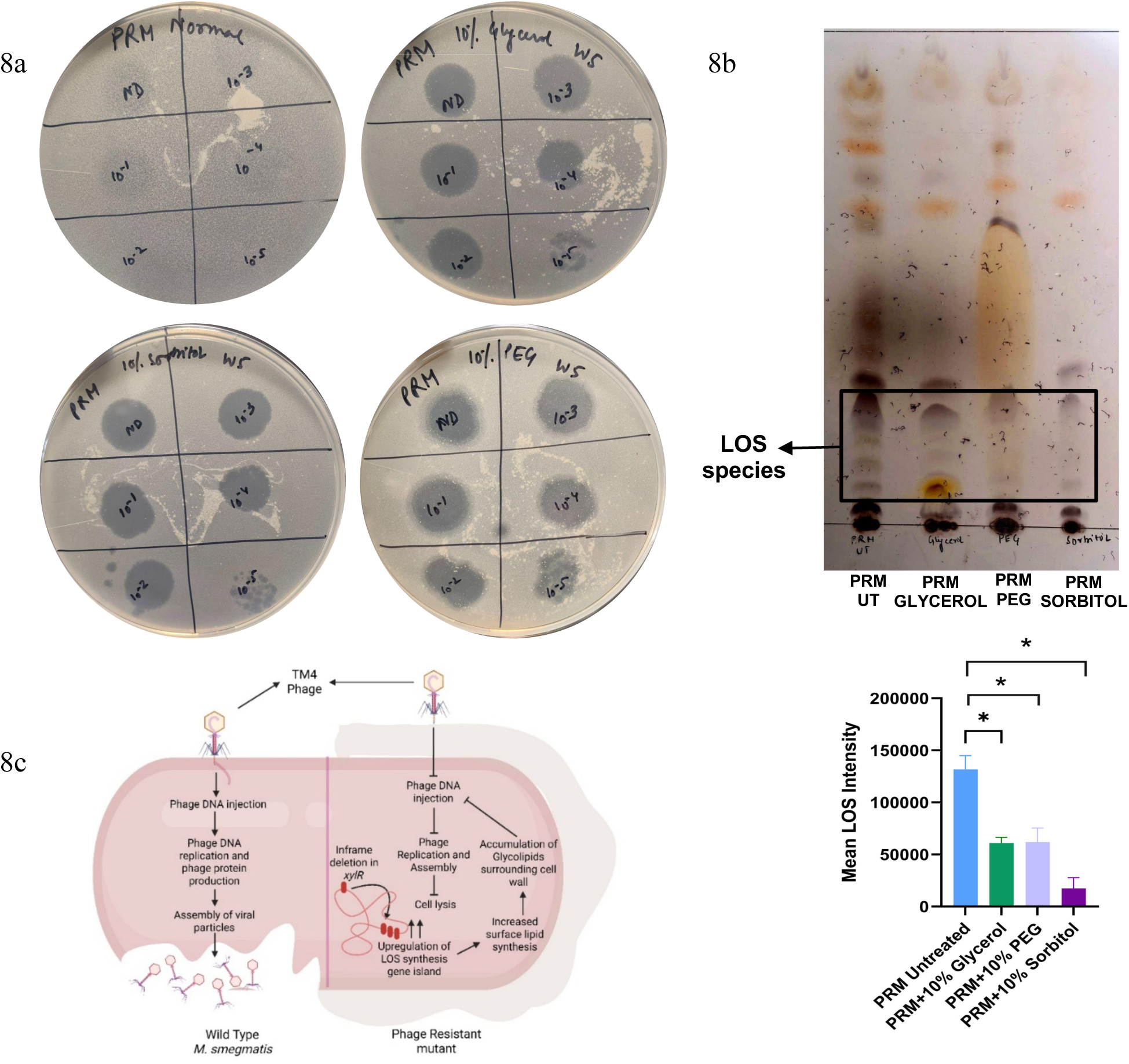
Loss of phage resistance in PRM with Polyol washes. **(a)** Polyol washes to PRM cells, reverses sensitivity to phage infection. Treatment was given with 10% Glycerol, 10% PEG, and 10% Sorbitol to the PRM cells. The experiment was replicated for at least 3 independent times. **(b)** Thin layer chromatography analysis of mycobacterial membrane lipids. TLC analysis of lipid extracts of untreated PRM, PRM washed with 10% glycerol, PRM washed with 10% PEG, and with 10% sorbitol in each case. Positions of LOS are indicated by arrows. Quantification of the LOS specific bands using image J. Statistical analysis was carried out using multiple Student’s *t*-tests. The *P*-values of the results (<0.05) are indicated by asterisk (*). **(c)** Disruption of *xylR* in *Mycobacterium smegmatis* leads to derepression of the LOS biosynthesis gene cluster, resulting in an increased production and accumulation of lipooligosaccharides at the cell surface. This accumulation creates a barrier around the cell that allows phage adsorption but inhibits the subsequent DNA injection into the host cell. Consequently, the infection cycle is arrested at an early stage, providing an effective, non-canonical defense mechanism against phage attack.

## DISCUSSION

Phage resistance in bacteria is commonly mediated by receptor modification, activation of antiviral systems (e.g., CRISPR-Cas, restriction–modification), or defective intracellular phage replication. Very few antiphage mechanisms have been clearly characterized in mycobacterial species, though bacterial genomes encode potential adaptive immune and innate immune systems^5–9^. Our phenotypic and transcriptomic data on the phage resistant mutant argue against these classical mechanisms. We observed that TM4 adsorption to resistant *M.smegmatis* cells remained unaffected, with no significant transcriptional changes observed in genes associated with known antiphage defense. However, the resistance phenotype is exhibited at an early post-adsorption stage, most plausibly during phage DNA injection. This suggests the role of cell envelope remodeling in providing an intrinsic structural barrier rather than a canonical immune defense.

An important finding of this study is the strong induction of the Lipooligosaccharide (LOS) biosynthesis gene cluster. This locus includes MSMEG_4727, the ortholog of the mycocerosic acid synthase like polyketide synthase *pks5*, responsible for generating polymethyl-branched fatty acids, key structural components of LOS. LOS-specific polar lipids species detected through TLC and MALDI-TOF MS confirmed their substantial accumulation in PRM, validating the transcriptomics analysis. Structural features of LOS, includes a poly-*O*-acylated trehalose core decorated with oligosaccharides and polymethyl-branched fatty acids. Enrichment of such polar lipids on the outer membrane, therefore, modifies the physicochemical properties of the cell envelope. Our data supports a model (Figure 8c) in which partial inactivation of the global regulator *xylR* leads to coordinated upregulation of the LOS biosynthesis gene island (MSMEG_4727–4737). This results in enhanced LOS accumulation on the cell surface that impedes TM4 phage to release its genome to the mycobacterial host to establish a successful infection. An alternate possibility suggests the binding of a bacteriophage to its host by a sequential process in which the phage makes multiple non-canonical interactions with the host cell surface. This interaction increases the contact time between the two entities thereby allowing the receptor binding domains in proteins subunits at the tail tip complex to bind tightly to the receptors. Indeed, a recent structural study have shown insights of the major capsid proteins of phage head to have glycan binding domains that can interact with the arabinose and mannose residues on the mycobacterial surface glycans thereby contributing in additional contacts with the bacterial surface^47^. We tested the susceptibility of a Δ*glgE* mutant strain that is deficient in producing the alpha glucan polysaccharides recruited to the mycobacterial outer surface. Indeed, TM4 phage showed a reduced adsorption and plaque formation on the Δ*glgE* mutant (Figure S3c, S3d)

The LOS specific gene locus (MSMEG_4727–4737) is repressed by *lsr2* and was earlier found to get upregulated upon *lsr2* inactivation. This loss of function also resulted in resistance towards mycobacteriophages^35^. Intriguingly, accumulation of mono-acylated species of the phosphatidylinositol mannoside and not LOS species were reported in the resistant mutant^35^. Here we present direct evidence of increased LOS deposition that alters the physicochemical properties of cell surface by increasing its hydrophilicity, in a way that it permits phage binding but prevents establishment of a successful infection due to the inability to deliver the phage genome. The augmented LOS layer may increase steric hindrance, modify membrane fluidity, or alter lipid organization at the phage–host interface. Such changes could disrupt the ability of mechanical penetration required for phage DNA injection. This mechanism is supported by our observation that phage genome replication and virion assembly were absent in PRM despite no defects in phage adsorption. Intriguingly, the observed resistant phenotype is reversible. Treatment of the resistant cells with polyols such as glycerol, sorbitol, or poly-ethelene glycol restored phage sensitivity and concomitantly reduced the accumulation of excess LOS species. This reversibility in phenotype shows that LOS accumulation leads to an adaptive surface modification rather than a permanent alteration in the core structure of the cell envelope. The ability to chemically modulate resistance further underscores the presence of a broadly specific induced innate immune response in Mycobacteria which results in an altered envelope rather than irreversible changes in phage receptors. Another possible reason for observing the insufficiency of phenotypic resistance at higher MOI which further diminished upon treatment with polyols could be due to a presence of a population heterogeneity wherein the cell wall of a sub-population did not show the accumulation of excessive LOS species on the surface. Microfluidics based single cell studies may help in mapping the extent of the non-resistors.

Our study establishes a clear link between *xylR* regulation and LOS overproduction to phage resistance, however, several important questions remain that requires further investigation. Direct experimental validation of a perturbed DNA injection process would strengthen our model, through experiments like fluorescent tracking of phage genome entry, or high-resolution imaging with cryo-EM to visualize the spatial and temporal progress of the TM4 phage–bacteria interactions. It would help to understand the nature of the trigger for phage genome delivery and the mechanism of the genome ejection against the positive hydrostatic pressure that the bacterial cytoplasm imparts on the phage head. Though, high resolution structural snapshots of Bxb1 phage infecting mycobacteria from cryo-EM studies have provided significant insights in our understanding of the dynamic interaction of the proteins of the phage tail-tip complex and the tail-spike with the mycobacterial outer and inner membrane^48^. This study also falls short in determining if the regulation of LOS gene cluster by *xylR* is through direct interaction, or via other regulatory proteins. Given the elevated metabolic activity observed in PRM, metabolomic and flux analyses could reveal how metabolic reprogramming supports enhanced lipid biosynthesis or can an increased oxidative stress response as we have observed constitutes a provides another broad-spectrum mechanism for phenotypic resistance.

Our observations also extend the functional scope of *xylR* beyond its classical role in xylose metabolism. Through this study we show that the LOS gene cluster is not only regulated by *lsr2*^35^, but is under the negative regulation of *xylR*. *xylR* acts as a central transcriptional regulator linking carbon and lipid metabolism and in turn regulating cell envelope structure. Inactivation of *xylR* de-represses the LOS biosynthetic genes, thereby conferring phage resistance. This reveals a novel signaling pathway connecting lipid metabolism and anti-phage defense and provides a new insight into host–phage co-evolution. Our findings highlight a broader implication for us to comprehend that cell envelope compositional plasticity-associated resistance mechanisms against phages representing a range of mycobacteriophage representing different clusters (A2, K2, B1) may develop a new manifestation of phenotypic antimicrobial resistance. This will eventually prove to become a major impediment in our effort towards optimizing phage-based therapeutic strategies against mycobacterial infections. Identifying unique phage resistance mechanisms in pathogenic mycobacteria would provide insights into the physiological and therapeutic potential of using phages and help us to design a phage cocktail that will be effective against antibiotic resistant strains of pathogenic Mycobacteria.

## MATERIALS AND METHODS

### Bacterial strains, phage strains, and growth conditions

*Mycobacterium smegmatis* mc²155 was cultured in Middlebrook 7H9 broth (BD Difco) supplemented with 0.2% glycerol and 0.05% Tween-80 at 37 °C with shaking at 180 rpm, or on Middlebrook 7H10 agar (BD Difco) supplemented with 0.2% glycerol. Phage-resistant *M. smegmatis* (PRM) cells were made electrocompetent and electroporated using a Bio-Rad Gene Pulser (25 µF, 2,500 V, 1000 Ω), and transformants were selected on 7H10 agar containing 20 µg/ml kanamycin; the constructs used in this study expressed the respective protein constitutively under the *hsp60* promoter. For phage infection assays, bacterial cultures were supplemented with 2 mM CaCl₂ and washed twice with MP buffer (50 mM Tris-HCl, pH 6.8; 150 mM NaCl; 1 mM MgSO₄; 2 mM CaCl₂) prior to infection. For phage propagation, wild-type Msm was grown on 7H10 plates at respective temperatures until confluency, after which the plates were flooded with 3 ml of MP buffer and incubated overnight at 4 °C with gentle shaking. The phage-containing buffer was collected, passed through a 0.22 µm filter to remove bacterial debris, and the resulting filtrate was used as the phage stock for subsequent experiments.

### Generation of spontaneous phage-resistant mutants and confirmation of phage resistance phenotype

5 ml cultures of 4 × 10^8^ colony-forming units (c.f.u.) *M. smegmatis* cells were incubated with MycoMar T7 phage lysate containing 2 × 10^9^ p.f.u. Phage (MOI 5) was incubated in a water bath at 30 °C with intermittent shaking for 25 hrs. Subsequently, 100 μl aliquots were then spread on Middlebrook 7H10 solid media after 3, 6, 9, 22, and 25 hrs, and incubated at 30 °C for 5 days or until isolated phage-resistant colonies were visible. Phage-resistant colonies were purified by streaking twice on Middlebrook 7H10 and, subsequently, were used to inoculate 5 ml cultures of Middlebrook 7H9 with 0.2% glycerol and 0.05% Tween 80. After growth to saturation, the culture was used to prepare bacterial lawns and serial dilutions of phages were spotted onto the lawns to determine phage susceptibilities.

### Extraction of DNA and performing Whole Genome Sequencing of the bacterial strains

Genomic DNA was isolated from both the parent *M. smegmatis* cells and the phage-resistant mutant following a previously described protocol^49^. Briefly, 20 mL of bacterial culture was harvested and resuspended in 500 µL of TE buffer (10 mM Tris-HCl, pH 8.0; 1 mM EDTA). To this suspension, 50 µL of lysozyme (10 mg/mL) was added, and the mixture was incubated overnight at 37 °C for complete cell wall digestion. Subsequently, 70 µL of 10% SDS and 50 µL of Proteinase K (10 mg/mL) were added and incubated at 60 °C for 1 h with shaking. Following lysis, 100 µL of pre-warmed 5 M NaCl and 10% CTAB were added to the mixture, which was then incubated at 60 °C for 15 min, frozen at −80 °C for 15 min, and again incubated at 60 °C for 15 min. The lysate was then extracted with chloroform–isoamyl alcohol, and the aqueous phase was carefully recovered. Genomic DNA was precipitated by adding ice-cold isopropanol and incubating at −80 °C for 2 h. The precipitated DNA was collected by centrifugation, washed with 70% ethanol, and air-dried. The purified DNA was finally dissolved in nuclease-free water, and its concentration and purity were assessed spectrophotometrically. Whole-genome sequencing of both the parent and mutant strains was performed using Sanger sequencing technology.

### Plasmid construction and complementation of the gene mutated in the phage-resistant mutant

To overexpress *xylR* in the phage-resistant mutant carrying a defective copy of the gene, the wild-type *xylR* was cloned into the mycobacterial expression vector pMV261. Briefly, the *xylR* gene was amplified from *M. smegmatis* genomic DNA using the primers 5′-GTGAGGAGT<u>GGATCC</u>GTTGGCAAACG-3′ (forward) and 5′

GAGCATCAA<u>AAGCTT</u>TCAACCGGC-3′ (reverse), which incorporated **BamHI** and **HindIII** restriction sites, respectively (sites underlined). PCR amplification was performed using Phusion High-Fidelity DNA Polymerase (New England Biolabs, USA) in the manufacturer-supplied buffer supplemented with 1.5 mM MgCl₂. The thermocycling conditions were as follows: initial denaturation at 95 °C for 3 min, followed by 35 cycles of 95 °C for 30 s, 56.2 °C for 30 s, and 72 °C for 1 min, with a final extension at 72 °C for 10 min.

The amplified *xylR* fragment was digested with **BamHI** and **HindIII** restriction enzymes, and ligated into the similarly digested pMV261 vector, generating the recombinant plasmid pMV261-*xylR*. The construct was confirmed by restriction digestion and sequencing. The resulting plasmid was electroporated into *M. smegmatis* mc²155 cells, which were subsequently recovered in MB7H9 broth supplemented with 0.2% glycerol and 0.05% Tween 80 (v/v) at 37 °C for 4 h. Following recovery, cells were plated onto MB7H10 agar containing 0.2% glycerol (v/v) and 20 µg/mL kanamycin for selection and incubated at 37 °C for 3 days.

### Phages and screening of phage susceptibility

Phages used in this study were obtained from the University of Pittsburgh, the University of Delhi, and the Indian Institute of Science. All the phages were propagated using *Mycobacterium smegmatis* mc²155 as the host strain. The lytic derivative phage ZoeJΔ43–45, a mutant of the temperate phage ZoeJ from Prof. Hatfull’s laboratory^40^, and phage 11 (PDRPxv), isolated from environmental samples in India by Prof. Urmi Bajpai’s group^41^, were included in the analysis.

Phage susceptibility of *M. smegmatis* strains was assessed using standard plaque assays. Top agar bacterial lawns were prepared by mixing 100 µl of mid-log-phase cell culture with molten Himedia top agar (0.6% BactoAgar, 2 mM CaCl₂) and overlaying onto MB7H10 agar plates. Tenfold serial dilutions of each phage were spotted onto the solidified lawns and incubated for 18–24 h at 37°C. Plaques were observed on the parental *M. smegmatis* strain, whereas the mutant strain displayed either complete resistance or reduced plaque-forming efficiency as compared to wild-type *M. smegmatis*, indicating a variable phage susceptibility.

### Morphological characterizations and determination of Mycobacterial growth curves

All *Mycobacterium smegmatis* strains were cultivated in Middlebrook 7H9 medium supplemented with 0.2% glycerol and 0.05% Tween 80 at 37 °C with continuous shaking at 180 rpm. Cultures were grown to mid-logarithmic phase (OD₆₀₀ ≈ 0.8), washed once with PBS-Tween 80 to remove any remaining media components. Approximately 20 cells were plated on Middlebrook 7H10 agar containing 0.2% glycerol to determine the colony morphology. Following incubation at 37 °C for 3–4 days, well-isolated colonies were visible, which were photographed using a digital camera.

To compare the growth curves of the *M. smegmatis* strains, cultures were seeded at an initial inoculum density of approximately 3.1 × 10⁴ cells ml⁻¹ (OD₆₀₀ ≈ 0.0003) in 50 ml of Middlebrook 7H9 medium supplemented with 0.2% glycerol and 0.05% Tween 80. The cultures were incubated at 37 °C with continuous shaking at 180 rpm. OD_600_ was recorded every 3 h to monitor bacterial growth until the cultures reached the stationary phase.

### Formation and Quantification of Mycobacterial Biofilms

To examine biofilm formation capacity, we followed previously described protocol with some modifications^50^. Briefly, *M. smegmatis* strains were first cultured in Middlebrook 7H9 medium supplemented with 0.2% glycerol and 0.05% Tween 80 at 37°C with shaking at 180 rpm until mid-logarithmic phase (OD₆₀₀ ≈ 0.8). The cultures were subsequently used to inoculate secondary cultures in the same medium. Log-phase cells were harvested and washed twice with M63 minimal medium (1 mM MgSO₄, 1 mM CaCl₂, casamino acids, (NH₄)₂SO₄, KH₂PO₄, FeSO₄·7H₂O, and 2% glycerol) to remove any residual media and Tween 80. The washed cells were resuspended in M63 medium and adjusted to an OD₆₀₀ of 0.2. 50 µl of these cell suspensions were seeded into each well of a sterile 24-well plate containing M63 medium, such that the number of cells per well corresponds to nearly 10⁶ CFU. The plate was incubated at 37°C undisturbed for 7 days inside a humidified chamber to allow for biofilm formation.

For quantification of biofilm biomass, the mature biofilms were stained with 2 ml of 0.1% (w/v) crystal violet for 10–15 min at room temperature. Excess stain was carefully removed, and the wells were washed 2–3 times with sterile distilled water to remove any unbound dye. To this, 2 ml of 95% ethanol was added to each well, followed by incubation for 10–20 min. A 125 µl aliquot of the resulting crystal violet–ethanol solution from each well was transferred to a 96-well flat-bottom plate, and the absorbance was measured at 630 nm using a SpectraMax iD3 plate reader.

### Ethidium Bromide uptake assay

To evaluate differences in cell wall permeability between strains, an ethidium bromide (EtBr) uptake assay was performed using a modified version of a previously described protocol^51^. Briefly, *M. smegmatis* cultures were grown in detergent-free Middlebrook 7H9 medium supplemented with 0.2% glycerol to mid-logarithmic phase (OD₆₀₀ ≈ 0.8). The cells were harvested, washed once with PBS, and subsequently pretreated with 0.4% glucose to energise the cells prior to EtBr uptake. The cell suspensions were then seeded into the wells of a black, clear-bottom 96-well plate containing EtBr prepared in PBS to final concentrations of 2 μg/mL. Fluorescence intensity was recorded at excitation/emission wavelengths of 530/590 nm at 1-minute intervals for 60 minutes using a SpectraMax iD3 plate reader.

### Measurement of Hydrophilicity Index

To determine the cell surface hydrophilicity of the *M. smegmatis* strains, a hexadecane partitioning assay was performed as previously described^52^. Cultures were grown in Middlebrook 7H9 media supplemented with 0.2% glycerol and 0.05% Tween 80 to mid-logarithmic phase (OD₆₀₀ ≈ 0.8). The cells were harvested and resuspended in PBS to a final OD₆₀₀ of 1.0 (OD_initial_). To each cell suspension, 1 ml of hexadecane was added, and the mixture was mixed vigorously to allow the mixing of both phases. The tubes were then allowed to stand undisturbed until the separation of the 2 phases occurred. The OD_600_ of the aqueous phase was measured (OD_final_). The hydrophilicity index (H-index) was calculated using the formula: H-index = (OD_final_/OD_initial_) X 100. A higher H-index indicates a more hydrophilic cell surface, whereas a lower H-index reflects greater hydrophobicity of the bacterial surface.

### Scanning electron microscopy

For scanning electron microscopy the procedure was adapted as described previously^53^. Briefly, the bacterial cell pellet obtained from a log-phase culture was fixed in the required buffer, attached to stubs, sputter coated with gold, and imaged on an UltraPlus scanning electron microscope (Zeiss, Germany) at 5.0 kV.

### MIC_99_ determination of antimicrobials and till-kill kinetics

The minimum inhibitory concentration (MIC) assay was performed in a 96-well plate with antimicrobials belonging to different classes: streptomycin, kanamycin, isoniazid, norfloxacin, rifampicin, ethambutol, D-cycloserine, and moxifloxacin. A serial 2-fold dilution was performed to obtain MIC_99_ values. Cells grown to the exponential phase (OD_600 nm_ ∼ 0.5) were diluted 1000 times to obtain OD_600 nm_ 0.0005 and seeded in detergent-free medium. A cell-free control and drug-free controls were maintained. Plates were incubated at 37 °C for 36 hrs before adding Resazurin at a final concentration of 30 μg/mL. After 5 hrs, fluorescence readings were taken at 530/590 nm using a Spectramax iD3 plate reader. For the Time-kill kinetics, cells with 0.8 OD_600nm_/mL (∼8 × 10^7^ CFU/mL) were challenged with different Kanamycin concentrations (fold MIC) at 37 °C at 200 RPM shaking. At regular intervals, cells were aliquoted and diluted in 7H9+0.05% Tween-80 and plated on LB agar to enumerate CFUs. After 3–4 days, the colonies were counted manually and plotted as CFU/mL over the course of time to exhibit a time-kill curve.

### Molecular modelling and dynamics simulation study of Wild type and mutant XylR

The domain and motif orientation of the wild type protein was carried out using MOTIF Search tool and Pfam domain analysis^54^. The secondary structure elements were predicted from FASTA amino acid sequence using the DSSP algorithm to determine the distribution of α-helices, β-sheets, and loop regions^55,56^. Since experimental structure of the XylR protein has not been determined yet, a homology modelling approach was adapted to generate the 3D structure of the wild-type XylR protein. Here in Swiss Modeler was used to determine the structure of the XylR protein^57^. The computationally determined XylR structure was verified for the stereochemical accuracy using Ramachandran plot. Additionally, the model was cross-verified segment by segment with the DSSP results to confirm the accuracy of the modelled structure. After successful construction and validation of the wild type XylR protein, the LVHG (AA162–165) deletion mutation was introduced to generate the mutant XylR. To investigate the conformational and dynamic variations of both the wild type and mutant proteins, molecular dynamics simulation was carried out for a period of 100 ns using GROMACS 2024 software package^58^. The GROMOS96 54a7 force field was utilized for generating the topology file for both wild type and mutated proteins for performing all atom molecular dynamics simulations. Before proceeding for the simulation, the systems were solvated in SPC water model inside a cubic box at 10 Å distance from the boundary of the protein structure. The systems were neutralized by adding appropriate counter ions to achieve a neutral net charge at physiological pH. To, eliminate the systems were energy minimized using both steepest decent and conjugate gradient methods for 50,000 steps. Before proceeding for the production MD simulation, the systems were subjected for two step equilibrations. The preliminary equilibration was carried out at constat temperature (300k) and further step of equilibration was carried out at constant pressure (1 bar). The equilibrated systems finally processed for the final step of 100ns MD simulation. The simulation trajectories were analyzed and compared to gain the critical confirmation changes.

### Growth curve of different Mycobacterial strains in the presence of TM4 Phage

To examine the growth curve of different *M. smegmatis* strains, they were first cultured in Middlebrook 7H9 medium supplemented with 0.2% glycerol and 0.05% Tween 80 at 37°C with shaking at 180 rpm until mid-logarithmic phase (OD₆₀₀ ≈ 0.8). The cultures were subsequently used to inoculate secondary cultures in the same medium. Log-phase cells were harvested and washed twice with MP Buffer to remove any residual media and Tween 80. The washed cells were resuspended in MP Buffer supplemented with 2mM CaCl₂, and adjusted to an OD₆₀₀ of 0.3. Aliquots of 100 µl (approximately 3 × 10⁷ CFU/ml) were seeded in the wells of a 96-well round-bottom microtiter plate. Each well was either infected with TM4 phage lysate at a multiplicity of infection (MOI) of 10 or received medium alone as an uninfected control. The plates were incubated at 37°C without agitation, and viable bacterial counts were determined every 3 hours until cultures reached the stationary phase.

### Adsorption assay of TM4 Phage in *M. smegmatis*

To find phage adsorption efficiency, *M. smegmatis* strains were grown to mid-logarithmic phase (OD₆₀₀ ≈ 0.8), washed twice with MP buffer and resuspended in MP buffer supplemented with 2 mM CaCl₂. Cells for the adsorption assay were taken such that the number of cells is approximately 10⁸ CFU/ml. TM4 phage lysate was added at an MOI of 0.01, and the samples were incubated in 30 ml glass tubes at 37°C with mild shaking (60 rpm) to aid phage adsorption. At different time points, 100 µl aliquots were collected, and the bacterial cells were pelleted by centrifugation. The supernatant, containing unadsorbed phages, was serially diluted and plated on lawns of wild-type *M. smegmatis* to determine the number of free phage particles. Plates were incubated at static 37°C for 21–24 hours until plaques appeared. All experiments were performed in biological triplicate.

### Generation of fluorescent phages with nucleic acid stain

Phages were stained with SYTOX™ Green nucleic acid stain as previously described^16^. Briefly, concentrated TM4 phage stocks (250 µL; 10¹⁰-10¹¹ p.f.u./mL) were incubated with SYTOX™ Green nucleic acid stain, at a final concentration of 0.5 µM for 30 mins at room temperature in the dark. Following incubation, the stained phages were washed four times with 2 mL of MP buffer using Amicon Ultra-15 centrifugal filter units to remove excess dye. After staining, the phage titre was immediately determined by plaque assay. Stained phage preparations were used within one week, as prolonged storage resulted in a gradual loss of viability.

### Flow cytometry

Phage adsorption efficiency was studied using a flow cytometry–based method as described previously^13,16^, with some modifications. Briefly, *M. smegmatis* strains were grown to mid-logarithmic phase (OD₆₀₀ ≈ 0.8), washed twice with MP buffer, and resuspended in 7H9 medium supplemented with 2 mM CaCl₂. The cell suspension was adjusted to an OD₆₀₀ of 0.1. The washed cells were then infected with SYTOX™ Green–stained TM4 phages at a multiplicity of infection (MOI) of 50 and incubated at 37 °C with gentle shaking (60 rpm) for 40 min. At 20 and 40 min post-infection, samples were transferred to ice to stop the phage infection process. The cells were then analysed by flow cytometry (Beckman Coulter CytoFLEX LX) at 488 nm excitation and 525 nm emission (B525-FITC-A). Bacterial populations were gated on SSC-A/FSC-A plots, and an unstained sample served as a negative control to define the SYTOX Green–positive gate. For each of the samples, more than 10,000 events were recorded. Data acquisition and analysis were performed using CytExpert software.

### Microscopic analysis of φ^2^GFP10-infected Mycobacterial strain

*Mycobacterium smegmatis* cultures were grown in 7H9 medium supplemented with 0.2% glycerol and 0.05% Tween-80 at 180 rpm till OD_600nm_ 0.8. After successive washes, bacterial cells were concentrated to obtain approximately 10⁷ CFU and subsequently infected with a fluorescently labelled variant of TM4 phage, φ2GFP10, at an MOI of 0.01. The infection was carried out at 30 °C with shaking at 60 rpm for 4 hours. Samples were collected at defined time points (0 h, 3 h, and 4 h post-infection) and imaged using confocal microscopy. Images were acquired in an Olympus FV3000 confocal microscope using a 100× oil immersion objective (NA 1.45). GFP fluorescence was excited with a 488 nm laser and detected at an emission wavelength of 514 nm. Images were captured using an ORCA Fusion sCMOS camera (5.3 MP) under identical acquisition settings.

### Image analysis

Representative Fields devoid of any artefacts were selected for analysis. Image processing was performed using Fiji software (fiji-windows-X64), with identical adjustments applied to intensity, brightness, and contrast between all the compared images sets.

### DNA electroporation and plaque quantification assay

The genomic DNA of TM4 phage was isolated using the phenol–chloroform extraction protocol as described previously^35^. Briefly, 500 ng of purified phage DNA was electroporated into different *Mycobacterium smegmatis* strains, followed by supplementation with 1 mL of 7H9 medium (without Tween-80). The cultures were then incubated at 37 °C with shaking at 180 rpm for 3 hours to allow for recovery. After recovery, the cells were harvested, resuspended in 200 µL of 7H9 medium, and mixed with 250 µL of wild-type *M. smegmatis*. The bacterial mixture was added to 3.5 mL of 0.6% molten agar and poured onto 7H10 agar plates. Plates were incubated at 37 °C, and plaques were counted after 24 hours.

### Sample preparations for RNASeq

Cells were grown in complete 7H9 medium till OD_600nm_ 0.8. After the cells were grown, they were harvested and washed with MP Buffer. To the washed cells, TM4 phage was added at an MOI of 10, and was incubated at 37 °C with shaking at 60 rpm for 140 minutes. Cells were collected at 0 minutes and 140 minutes post-infection and was harvested and washed with ice-cold PBS-Tween80, to stop any further phage infection. After washing, cells were freeze frozen using Liquid nitrogen and stored at -80 °C until further processing. RNA extraction was performed by using the Qiagen miRNeasy Mini kit (Cat. No. 1038703).

### Transcriptomic data analysis

The entire process of RNA extraction, quality assessment, library preparation and high-throughput sequencing were all outsourced to a commercial sequencing facility. Raw sequencing reads were processed, aligned to the *Mycobacterium smegmatis* reference genome, and analysed to identify differentially expressed genes (DEGs) between the various conditions. Genes exhibiting a log₂ fold change (Log₂FC) ≥ 1.5 and a *p*-value ≤ 0.05 were considered significant, and selected for downstream analyses. To understand the changes in gene expression profile in WT and PRM, the identified DEGs were functionally categorized based on their cellular pathways and physiological roles they are involved in. Functional classification was performed using categories analogous to those established for *Mycobacterium tuberculosis*. The DEGs were grouped into categories including intermediary metabolism and respiration, lipid metabolism, cell wall and cell processes, membrane transport, regulatory proteins, conserved hypotheticals, virulence-associated functions, information pathways, etc. Following functional catagorization, graphical representations to visualize the distribution and abundance of DEGs within each category were made, thereby facilitating identification of the major biological processes altered in PRM.

### Measurement of respiratory activity using Resazurin

Measurement of metabolic/respiratory activity was performed as previously reported^53^. Briefly, cells were grown till the exponential phase in complete 7H9 medium. At a certain time point, 100 µl aliquots containing equal numbers of cells were collected and transferred into black-walled 96-well plates (Costar). Resazurin was added to all wells at a final concentration of 30 µg/ml and incubated in the dark at 37 °C for 30 min. The quick conversion of Resazurin to resorufin was measured in a plate reader at 530/590 nm. Higher the conversion of Resazurin to resorufin, higher is the metabolic rate of that strain.

### Detection of basal levels of ROS in WT and PRM using CellROX Deep Red dye

Measurement of basal levels of ROS was performed as reported earlier^53^. Briefly, logarithmic phase WT and PRM cells were treated with 5 µM of CellROX Deep Red dye and 200µl was transferred in the dark black wall 96-well plate (Costar) and incubated at 37 °C for 40 min. Fluorescence was recorded at 645/665 nm in a micro plate reader (BioTek Synergy H1). Fluorescence intensities were divided by CFUs, measured just before the addition of the CellROX deep red dye to determine ROS levels per bacterium. CFU normalized fluorescence intensities for both WT and PRM were plotted in the graph.

### Extraction and purification of mycobacterial lipids

The total lipid fraction was extracted from cell pellets with a mixture of chloroform and methanol as previously described^59^. The various lipid extracts were dissolved in chloroform: methanol (2:1) and were analyzed by thin-layer chromatography (TLC) on silica gel Durasil 25-precoated plates (0.25-mm thickness; Macheray-Nagel). LOS were resolved by TLC run in chloroform-methanol (90:10, vol/vol) and visualized by spraying the plates with alpha napthol, followed by heating at 110°C. Purification of LOS was achieved by preparative TLC in which plates were developed as described above, and then LOS-containing silica bands were scraped and extracted three times with the same solvent.

### MALDI-TOF Mass Spectrometry

After running the TLC as per the above protocol, LOS specific regions on the TLC plate was scraped carefully. The scraped silica was collected in a glass vial and was resuspended in chloroform-methanol (90:10, vol/vol), and this stock was stored at 4°C for further use. For the sample preparation for MALDI-TOF, we used the same protocol as previously described^60^, as matrix solutions, 2,5-Dihydroxybenzoic acid (2,5-DHB) was used. MALDI-TOF mass spectra (in the positive ion mode) were acquired on a UTG-10132 RapifleX mass spectrometer (Bruker Daltonics, USA) equipped with a pulsed nitrogen laser emitting at 337 nm. Samples were analyzed in the Reflectron mode and an accelerating voltage operating in positive ion mode of 20 kV. An external mass spectrum calibration was performed using the calibration mixture 1 of Peptide Calibration Standard mono, including known peptide standards in a mass range from 900 to 1600 Da.

### Polyol washes of the PRM cells and performing plaque assays

PRM cells were grown in complete 7H9 medium until reaching an OD₆₀₀ of 0.8. Cultures were harvested at 4°C and washed three times with ice-cold 10% glycerol, 10% sorbitol, or 10% polyethylene glycol (PEG), followed by final resuspension in the respective solvent. The treated cells were subsequently used for plaque assays following the protocol described above.

### Statistical analysis

All experiments were performed at least three times or more independently. All the data are represented as mean, and the error bars represent the standard deviation. All statistical analyses were performed using GraphPad Prism 10. For the calculation of statistical significance, an unpaired, two-tailed Student’s t-test was performed in a pair-wise manner (between control and the desired experiment), as indicated in the figures.

## ACKNOWLEDGEMENT

AA and RS thanks UGC and DBT for graduate research fellowships respectively. RM thanks ICMR (Grant no: IRPSG-2024-01-01114), DBT (Grant no: BT/PR20820/MED/30/1875/2017) and IISER Tirupati for funding. Authors thank Drs. Bill Jacobs, Graham Hatfull, and Urmi Bajpai for providing φ^2^GFP10, D29, TM4, ZoeJΔ43-45 and PDRPxv phages, respectively, and Rainer Kalscheuer for providing the Δ*glgE* mutant strain. The authors also thank Dr. Vikas Jain for SEM analysis.

