## Supplementary figures for "Mycobacteriophage TM4 requires XylR for successful infection in *Mycobacterium smegmatis* mc^2^155"

### Supplementary information

S1a

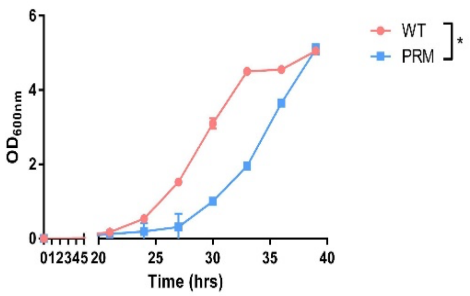


S1b

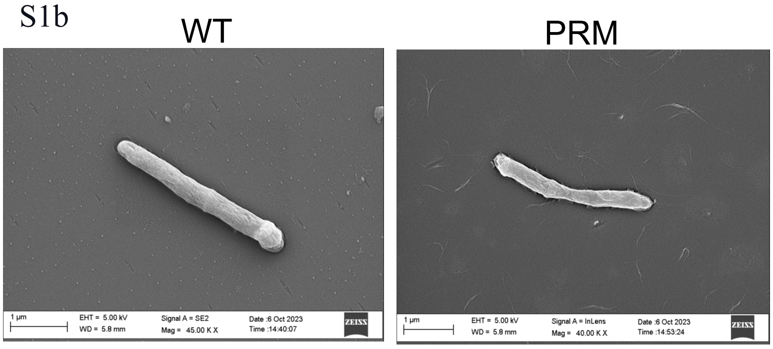


S1c

| | MINIMUM INHIBITORY CONCENTRATIONS (MIC99) (μg/mL) | | | | | | | |
|---|---|---|---|---|---|---|---|---|
| STRAINS \ ANTIBIOTICS | Streptomycin | Kanamycin | Isoniazid | Ethambutol | Rifampicin | Cycloserine | Norfloxacin | Moxifloxacin |
| WT | 0.125 | 2 | 2 | 0.25 | 0.3125 | 80 | 2 | 0.0625 |
| PRM | 0.125 | 1 | 4 | 0.25 | 0.3125 | 40 | 2 | 0.0625 |

S1d

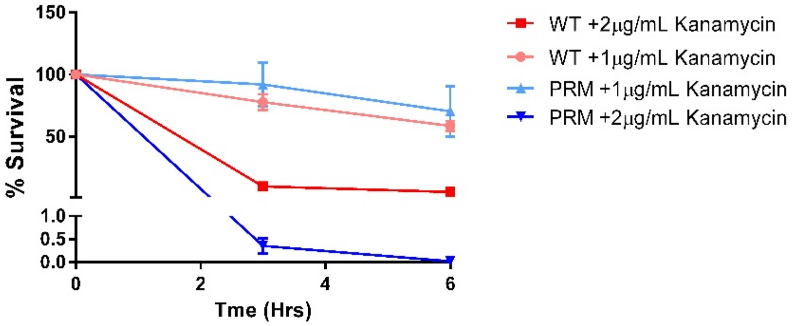


**Figure S1: Phenotypic alterations n PRM as compared to WT *M.smegmatis***

**(a)** Growth curve of WT *M.smegmatis* and PRM in presence of complete 7H9 medium

**(b)** SEM of WT *M.smegmatis* and PRM

**(c)** $MIC_{99}$ of different antimicrobials against WT and PRM cells using resazurin reduction assay.

**(d)** Calculating the CFU after treating WT and PRM with different concentrations of Kanamycin. All the tests were performed in three biological replicates, and error bars indicate the standard deviations. Statistical analysis was carried out using multiple Student's *t*-tests. The *P*-values of the results <0.05 is indicated by asterisk (*).

S2a

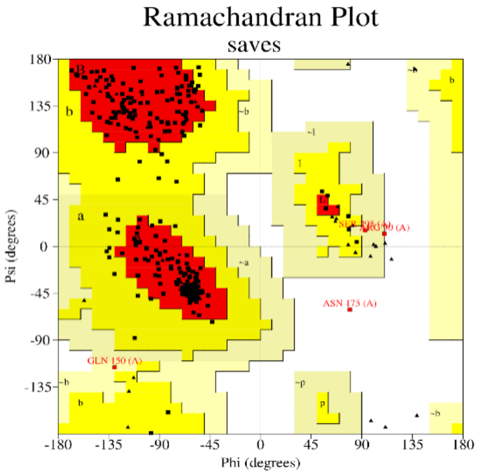


S2b

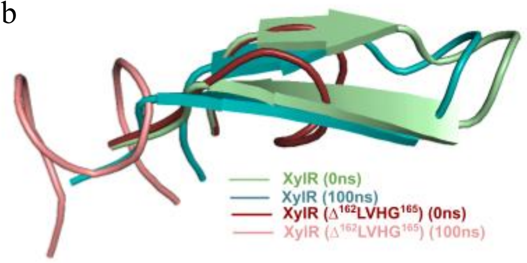


S2c

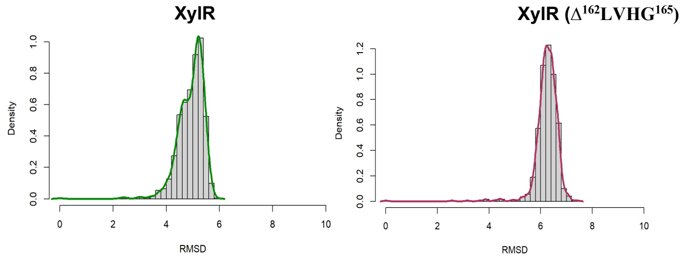


S2d

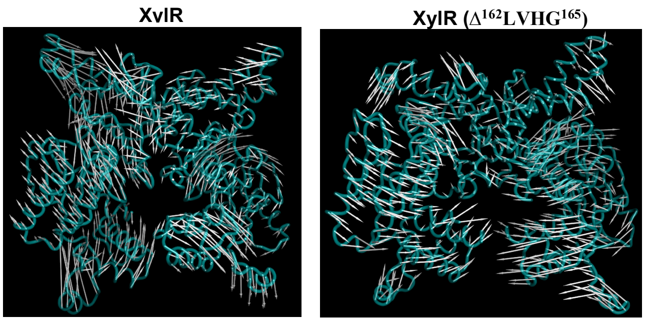


S2e

| XylR-DNA Complex | | | | XylR ($\Delta^{162}$LVHG$^{165}$)-DNA Complex | | | |
|---|---|---|---|---|---|---|---|
| HADDOCK score: -36.2 +/- 5.8<br>Z-Score: -2.4<br>Binding energy: -50.668 kcal/mol | | | | HADDOCK score: -11.7 +/- 6.0<br>Z-Score: -1.5<br>Binding energy: -46.616 kcal/mol | | | |
| **H-Bonds** | **Protein residues** | **DNA residues** | **Bond Length (Å)** | **H-Bonds** | **Protein residues** | **DNA residues** | **Bond Length (Å)** |
| **Chain A** | 51ARG | DA92 | 2.77 | **Chain A** | 39SER | DT23 | 2.64 |
| | 55LYS | DG91 | 2.99 | | 40ARG | DC24 | 2.71 |
| | 437SER | DG32 | 3.02 | | 41ALA | DC24 | 2.61 |

| | | | | | | | |
|---|---|---|---|---|---|---|---|
| **Chain B** | 438ARG | DA34 | 2.54 | | 51ARG | DG91 | 2.47 |
| | 438ARG | DT81 | 2.50 | | 53THR | DG89 | 3.04 |
| | 439SER | DG32 | 2.08 | | 56ASP | DG89 | 2.61 |
| | 440THR | DG32 | 2.10 | **Chain B** | No H-Bonds observed | | |
| | 440THR | DG32 | 2.30 | | | | |
| | 442LYS | DT84 | 3.05 | | | | |

**Figure S2: Comparative structural and dynamic analysis for XylR**

- **(a)** The Ramachandran plot showing the distribution of backbone dihedral angles (φ and ψ) of the wild type XylR. Red and yellow region corresponds to most favourable and allowed regions.
- **(b)** MD simulation showing the change in secondary structure at the site of mutation over the course of 100ns.
- **(c)** The probability density distribution of RMSD values XylR obtained from the MD trajectory.
- **(d)** The porcupine plots illustrating the change in overall domain movement of the mutant protein as compare to the WT XylR.
- **(e)** Table showing the atomic interactions (H-bonds) between the interacting partners of promoter DNA and XylR.

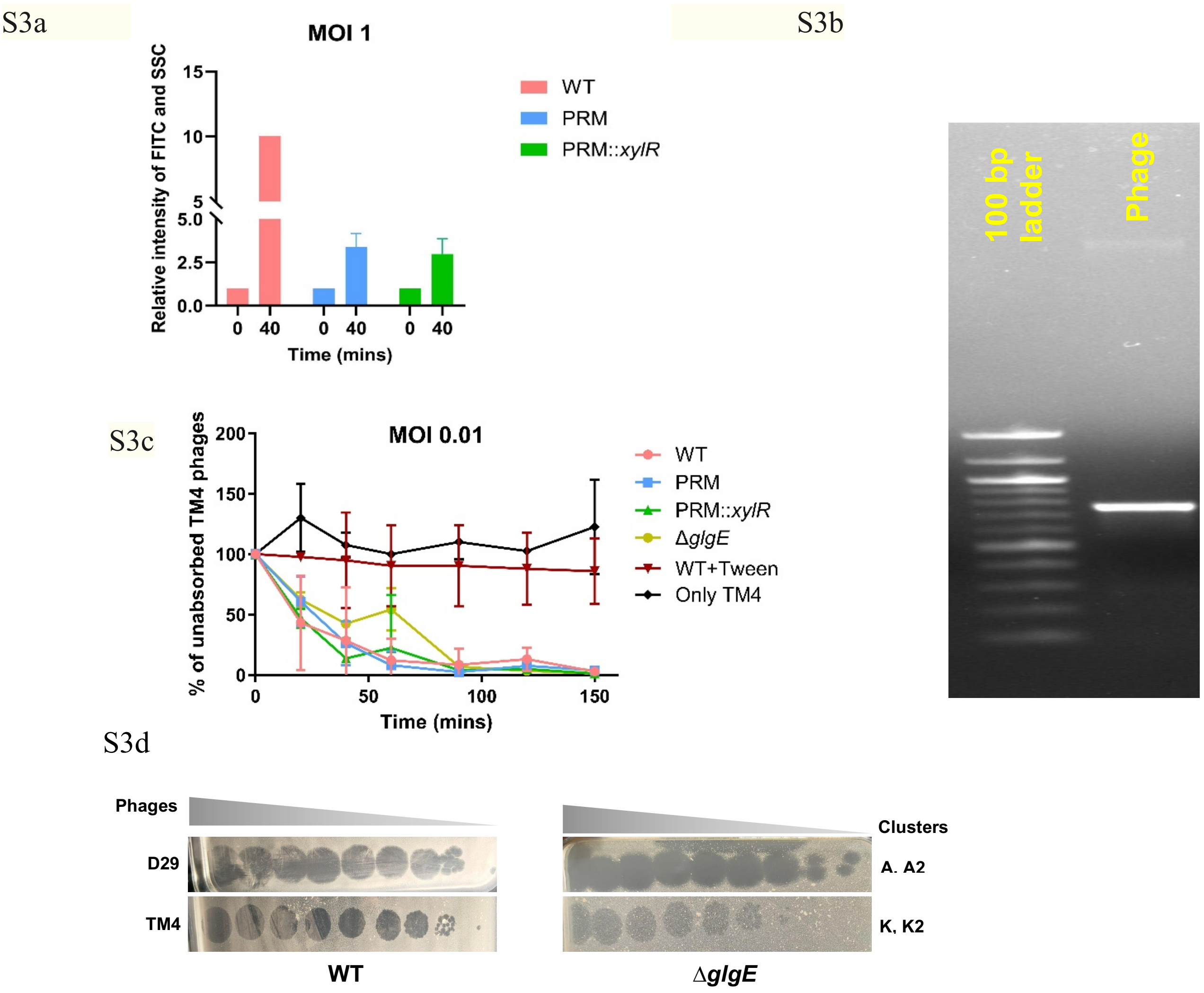


**Figure S3: TM4 Phage adsorption is independent of *xylR* regulation**

- **(a)** Graphical representation of the flow cytometry data. The number represents the events recorded in Q1 quadrant, recorded from cells positive for both SSC and FITC. Data is represented for all the 3 strains for 3 different time points. Increase in the Q1 population represents phage adhered bacterial cells.
- **(b)** Verification of isolated TM4 phage genomic DNA. TM4 gDNA was amplified using phage capsid specific (right panel) primers. 'Phage' represents TM4 phage DNA.
- **(c)** Comparison for adsorption of phage TM4 to different mycobacterial strains, including WT, PRM, PRM::*xylR*, or Δ*glgE* strain. All the tests were performed in three biological replicates, and error bars indicate the standard deviations.
- **(d)** Phage susceptibility of WT, and Δ*glgE* strains towards different Mycophages. Tenfold serial dilutions of phage lysates (D29, TM4) were spotted on solid media with bacterial lawns. Plaque assays were performed at least three independent times with consistent results.

S4

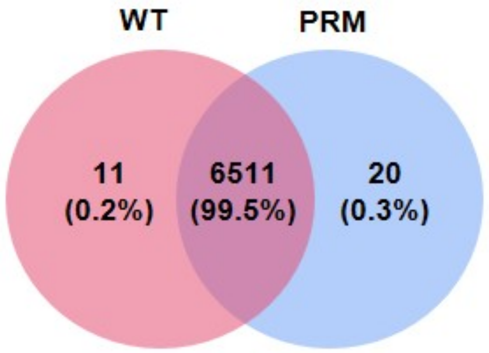


**Figure S4: Comparing gene expression profile between WT and PRM cells**

Venn diagram showing commonality between the WT and PRM untreated transcriptomic dataset. The overlap indicates the number of genes which are mutually present in both the samples.

S5a

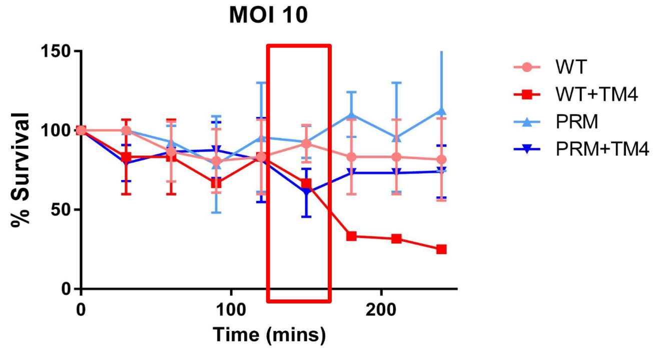


S5b

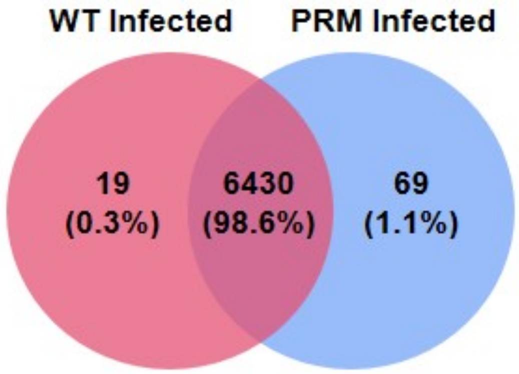


S5c

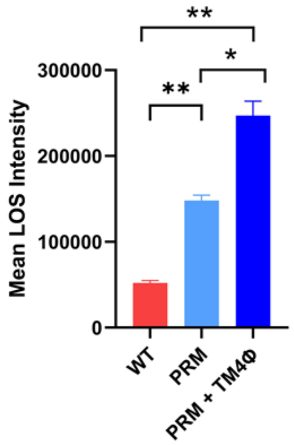


S5d

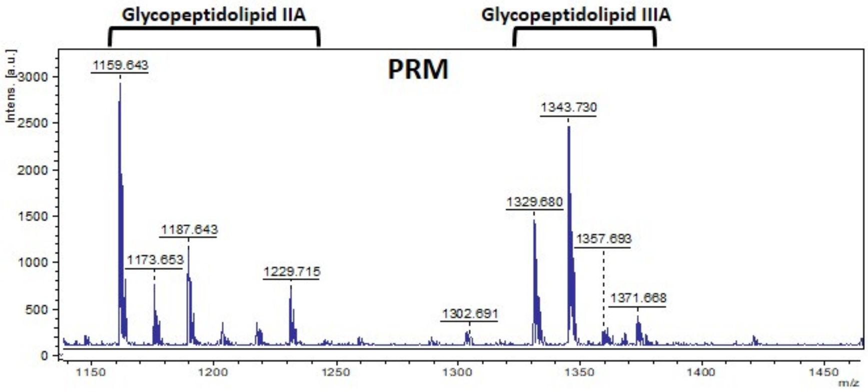


**Figure S5: Comparing the gene expression profile of WT and PRM cells upon TM4 phage infection**

**(a)** Growth curve of WT and PRM in presence of TM4 phage infection. Time point of 140 mins was selected for harvesting the cells for transcriptomics.

**(b)** Venn diagram showing commonality between the WT and PRM TM4 phage infected transcriptomics profile. The overlap indicates the number of genes which are mutually present in both the samples.

**(c)** Quantitation of the LOS specific bands using image J. Statistical analysis was carried out using multiple Student's *t*-tests. The *P*-values of the results ($<0.05$, $<0.01$) are indicated by asterisk (*, **).

**(d)** MALDI-TOF MS of the lipid extracts of the WT, and PRM strains. Major Pseudo molecular ion $[M + Na]^+$ peaks of GPLs are indicated with the spectra.
